# Modular metabolite assembly in *C. elegans* depends on carboxylesterases and formation of lysosome-related organelles

**DOI:** 10.1101/2020.08.22.262956

**Authors:** Henry H. Le, Chester J. J. Wrobel, Sarah M. Cohen, Jingfang Yu, Heenam Park, Maximilian J. Helf, Brian J. Curtis, Joseph C. Kruempel, Pedro R. Rodrigues, Patrick J. Hu, Paul W. Sternberg, Frank C. Schroeder

## Abstract

Signaling molecules derived from attachment of diverse metabolic building blocks to ascarosides play a central role in the life history of *C. elegans* and other nematodes; however, many aspects of their biogenesis remain unclear. Using comparative metabolomics, we show that a pathway mediating formation of intestinal lysosome-related organelles (LROs) is required for biosynthesis of most modular ascarosides as well as previously undescribed modular glucosides. Similar to modular ascarosides, the modular glucosides are derived from highly selective assembly of moieties from nucleoside, amino acid, neurotransmitter, and lipid metabolism, suggesting that modular glucosides, like the ascarosides, may serve signaling functions. We further show that carboxylesterases that localize to intestinal organelles are required for the assembly of both modular ascarosides and glucosides via ester and amide linkages. Further exploration of LRO function and carboxylesterase homologs in *C. elegans* and other animals may reveal additional new compound families and signaling paradigms.

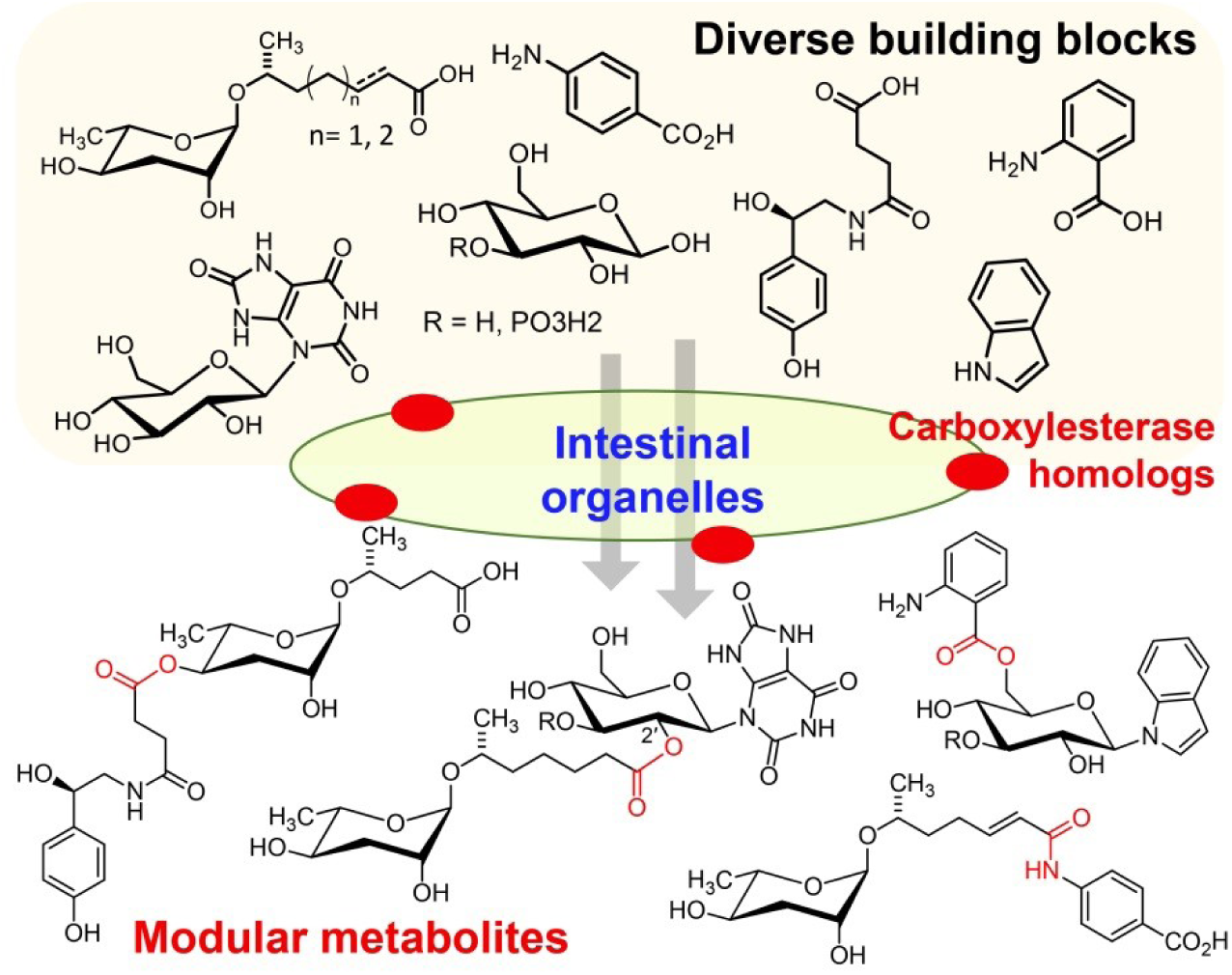

## Introduction

Recent studies indicate that the metabolomes of animals, from model systems such as *Caenorhabditis elegans* and Drosophila to humans, may include >100,000 of compounds^1, 2^. The structures and functions of most of these small molecules have not been identified, representing a largely untapped reservoir of chemical diversity and bioactivities. In *C. elegans*^3^ a large modular library of small-molecule signals, the ascarosides, are involved in almost every aspect of its life history, including aging, development, and behavior^4–7^. The ascarosides represent a structurally diverse chemical language, derived from glycosides of the dideoxysugar ascarylose and hydroxylated short-chain fatty acid (Fig. 1a)^8^. Structural and functional specificity arises from optional attachment of additional moieties to the sugar, for example indole-3- carboxylic acid (e.g. icas#3 (**1**)), or carboxy-terminal additions to the fatty acid chain, such as *p*- aminobenzoic acid (PABA, as in ascr#8 (**2**)) or *O*-glucosyl uric acid (e.g. uglas#11 (**3**), Fig. 1b)^2, 9–13^. Given that even small changes in the chemical structures of the ascarosides often result in starkly altered biological function, ascaroside biosynthesis appears to correspond to a carefully regulated encoding process in which biological state is translated into chemical structures^14^. Thus, the biosynthesis of ascarosides and other *C. elegans* signaling molecules (e.g. nacq#1)^15^ represents a fascinating model system for the endogenous regulation of inter- organismal small-molecule signaling in metazoans. However, for most of the >200 recently identified *C. elegans* metabolites^2, 8, 9^, biosynthetic knowledge is sparse. Previous studies have demonstrated that conserved metabolic pathways, e.g. peroxisomal *β*-oxidation^9, 10^ and amino acid catabolism^8, 16^ (Fig. 1a), contribute to ascaroside biosynthesis; however, many aspects of the mechanisms underlying assembly of multi-modular metabolites remains unclear.

**Figure 1:**
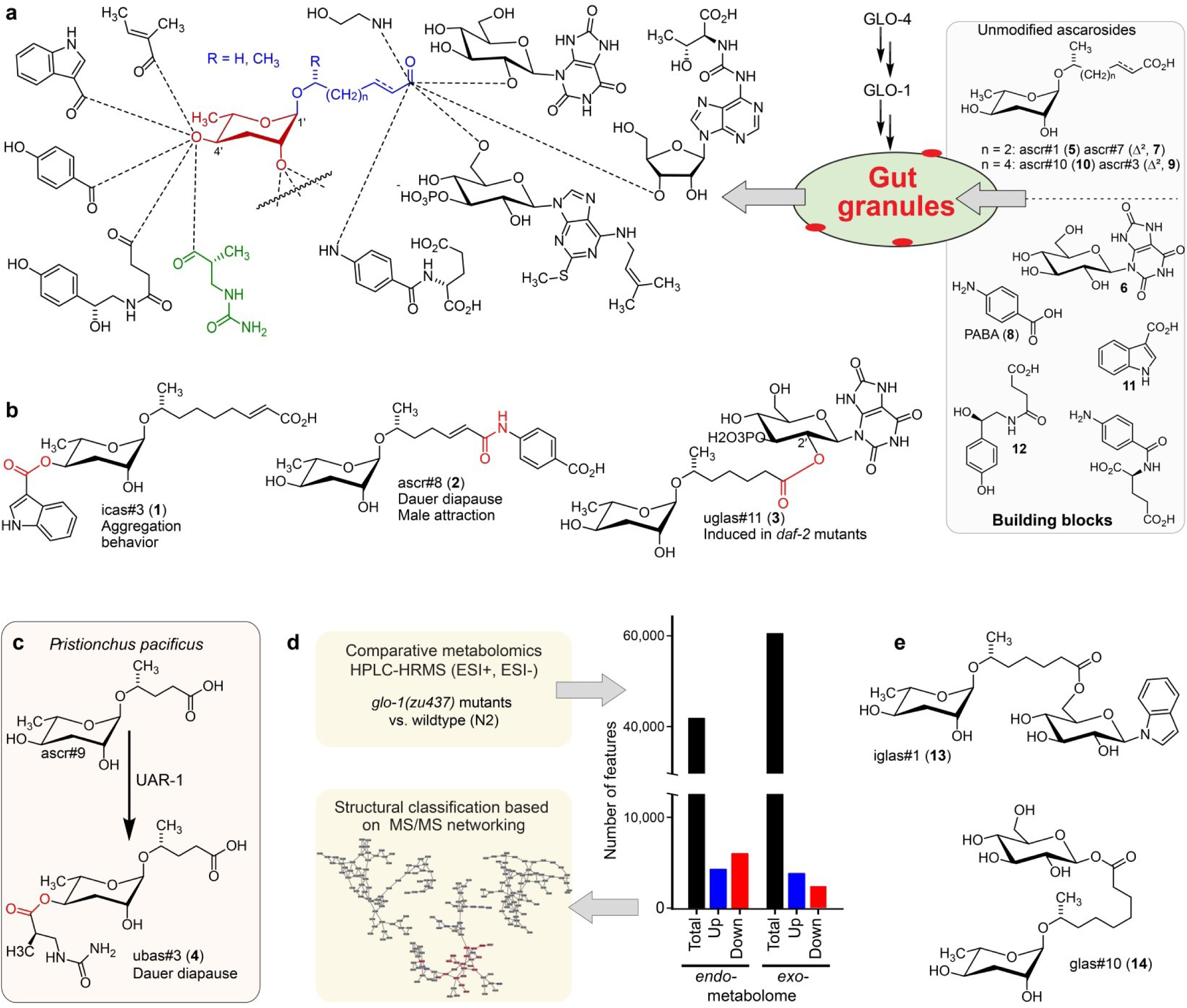
(a) Modular ascarosides are assembled from simple ascarosides, e.g. ascr#1 (**5**) or ascr#3 (**9**), and building blocks from other metabolic pathways, e.g. glucosyl uric acid (**6**), *p*- aminobenzoic acid (PABA, **8**) indole-3-carboxylic acid (**11**), or succinyl octopamine (**12**). We hypothesize that *glo-1*-dependent gut granules play a central role in their biosynthesis. (b) Examples for modular ascarosides and their biological context. (c) UAR-1 in *P. pacificus* converts simple ascarosides into the 4′-ureidoisobutyric acid-bearing ascarosides, e.g. ubas#3 (**4**). (d) Strategy for comparative metabolomic analysis of LRO-deficient *glo-1* mutants. (e) Example for modular ascarosides whose production is increased in *glo-1* mutants.

Recently, metabolomic analysis of mutants of the Rab-GTPase *glo-1*, which lack a specific type of lysosome-related organelles (LROs, also referred to as autofluorescent gut granules), revealed complete loss of 4′-modified ascarosides^14^. The *glo-1*-dependent LROs are acidic, pigmented compartments that are related to mammalian melanosomes and drosophila eye pigment organelles^17, 18^. LROs form when lysosomes fuse with other cellular compartments, e.g. peroxisomes, and appear to play an important role for recycling proteins and metabolites^17^. Additionally, it has been suggested that LROs may be involved in the production and secretion of diverse signaling molecules^19, 20^, and the observation that *glo-1* mutant worms are deficient in 4′-modified ascarosides suggested that intestinal organelles may serve as hubs for their assembly (Fig. 1a)^14^. In addition to the autofluorescent LROs, several other types of intestinal granules have been characterized in *C. elegans*, including lipid droplets^21^ and lysosome-related organelles that are not *glo-1*-dependent ^22^.

Parallel studies of other *Caenorhabditis* species^23–25^ and *Pristionchus pacificus*^26^, a nematode species being developed as a satellite model system to *C. elegans*^27^, revealed that production of modular ascarosides is widely conserved among nematodes. Leveraging the high genomic diversity of sequenced *P. pacificus* isolates, genome-wide association studies coupled to metabolomic analysis revealed that *uar-1*, a carboxylesterase from the α/ß-hydrolase superfamily with homology to cholinesterases (AChEs), is required for 4′-attachment of an ureidoisobutyryl moiety to a subset of ascarosides, e.g. ubas#3 (**4**, Fig. 1c)^26^. Homology searches revealed a large expansion of <u>c</u>arboxyl<u>est</u>erase (*cest*) homologs in *P. pacificus* as well as *C. elegans* (Fig. S1), and recently it was shown that in *C. elegans*, the *uar-1* homologs *cest- 3*, *cest-8*, and *cest-9.2* are involved in the 4′-attachment of other acyl groups in modular ascarosides^28^. Based on these findings, we posited that *cest* homologs localize to *glo-1*- dependent intestinal granules where they control assembly of modular ascarosides, and perhaps other modular metabolites. In this work, we present a comprehensive assessment of the impact of *glo-1*-deletion on the *C. elegans* metabolome and uncover the central role of *cest* homologs that localize to intestinal granules in the biosynthesis of diverse modular metabolites.

## Results

### Novel classes of LRO-dependent metabolites

To gain a comprehensive overview of the role of *glo-1* in *C. elegans* metabolism, we employed a fully untargeted comparison of the metabolomes of a *glo-1* null mutant and wildtype worms (Fig. 1d). HPLC-high resolution mass spectrometry (HPLC-HRMS) data for the *exo*-metabolomes (excreted compounds) and *endo-* metabolomes (compounds extractable from the worm bodies) of the two strains were analyzed using the Metaboseek comparative metabolomics platform, which integrates the *xcms* package^29^. These comparative analyses revealed that the *glo-1* mutation has a dramatic impact on *C. elegans* metabolism. For example, in negative ionization mode we detected >1000 molecular features that were at least 10-fold less abundant in the *glo-1 exo-* and *endo*- metabolomes, as well as >3000 molecular features that are 10-fold upregulated in *glo-1* mutants. For further characterization of differential features, we employed tandem mass spectrometry (MS^2^) based molecular networking, a method which groups metabolites based on shared fragmentation patterns (Fig. 1d, S2-5)^30^. The resulting four MS^2^ networks – for data obtained in positive and negative ionization mode for the *exo-* and *endo-*metabolomes – revealed several large clusters of features whose abundances were largely abolished or greatly increased in *glo-1* worms. Notably, although some differential MS^2^ clusters represented known compounds, e.g. ascarosides, the majority of clusters were found to represent previously undescribed metabolite families.

In agreement with previous studies^14^, biosynthesis of most modular ascarosides was abolished or substantially reduced in *glo-1* mutants, including all 4′-modified ascarosides, e.g. icas#3 (**1**) (Figs. 1b, and S6a). Similarly, production of ascarosides modified at the carboxy terminus, e.g. uglas#11 (**3**) derived from ester formation between ascr#1 (**5**) and uric acid glucoside^12^ (**6**), and ascr#8 (**2**), derived from formation of an amide bond between ascr#7 (**7**) and of *p*-amino benzoic acid (**8**), was largely abolished in *glo-1* mutants (Figs. 1a, 1b, and S6a). Metabolites plausibly representing building blocks of these modular ascarosides were not strongly perturbed in *glo-1* mutants (Fig. S7). For example, abundances of unmodified ascarosides, e.g. ascr#3 (**9**) and ascr#10 (**10**), or metabolites representing 4′-modifications, e.g. indole-3-carboxylic acid (**11**) and octopamine succinate (**12**), were not significantly perturbed in the mutant (Figs. 1a, S6a and S7). In contrast, a subset of modular ascaroside glucose esters (e.g. iglas#1 (**13**) and glas#10 (**14**), Fig. 1e), was strongly increased in *glo-1* mutants (Fig. S6b). These results suggest that *glo-1*-dependent intestinal organelles function as a central hub for the biosynthesis of most modular ascarosides, with the exception of a subset of ascarosylated glucosides, whose increased production in *glo-1* mutants may be indicative of a shunt pathway for ascarosyl-CoA derivatives^31–33^, which represent plausible precursors for modular ascarosides modified at the carboxy terminus.

Next, we analyzed the most prominent MS^2^ clusters representing previously uncharacterized metabolites whose production is abolished or strongly reduced in *glo-1* mutants (Fig. 2). Detailed analysis of their MS^2^ spectra indicated that they may represent a large family of modular hexose derivatives incorporating moieties from diverse primary metabolic pathways. For example, MS^2^ spectra from clusters **I**, **II**, and **III** of the positive-ionization network suggested phosphorylated hexose glycosides of indole, anthranilic acid, tyramine, or octopamine, which are further decorated with a wide variety of fatty acyl moieties derived from fatty acid or amino acid metabolism, for example nicotinic acid, pyrrolic acid, or tiglic acid (Fig. 2, Table 1)^17, 34^. Given the previous identification of the glucosides iglu#1/2 (**15/16**, Fig. 2e) and angl#1/2 (**17/18**), we hypothesized that clusters **I**, **II**, and **III** represent a modular library of glucosides, in which *N*-glucosylated indole, anthranilic acid, tyramine, or octopamine^35^ serve as scaffolds for attachment of diverse building blocks. To further support these structural assignments, a series of modular metabolites based on *N*-glucosylated indole (“iglu”) were selected for total synthesis. Synthetic standards for the non-phosphorylated parent compounds of iglu#4 (**19**), iglu#6 (**20**), iglu#8 (**21**), and iglu#10 (**22**) matched HPLC retention times and MS^2^ spectra of the corresponding natural compounds (Fig. S8), confirming their structures and enabling tentative structural assignments for a large number of additional modular glucosides, including their phosphorylated derivatives, e.g. iglu#12 (**23**), iglu#41 (**24**), angl#4 (cluster **II**, **25**), and tyglu#4 (cluster **III**, **26**) (Fig. 2). The proposed structures include several glucosides of the neurotransmitters tyramine and octopamine, whose incorporation could be verified by comparison with data from a recently described feeding experiment with stable isotope labeled tyrosine^35^. Similar to ascaroside biosynthesis, the production of modular glucosides is life stage dependent; for example, production of specific tyramine glucosides peaks at the L3 larval stage, whereas production of angl#4 increases until the adult stage (Figs. S9 and S10). Notably, modular glucosides were detected primarily as their phosphorylated derivatives, as respective non-phosphorylated species were generally less abundant. In contrast to most ascarosides, the phosphorylated glucosides are more abundant in the *endo*-metabolome than the *exo*- metabolome, suggesting that phosphorylated glucosides may be specifically retained in the body (Fig. S9).

**Figure 2:**
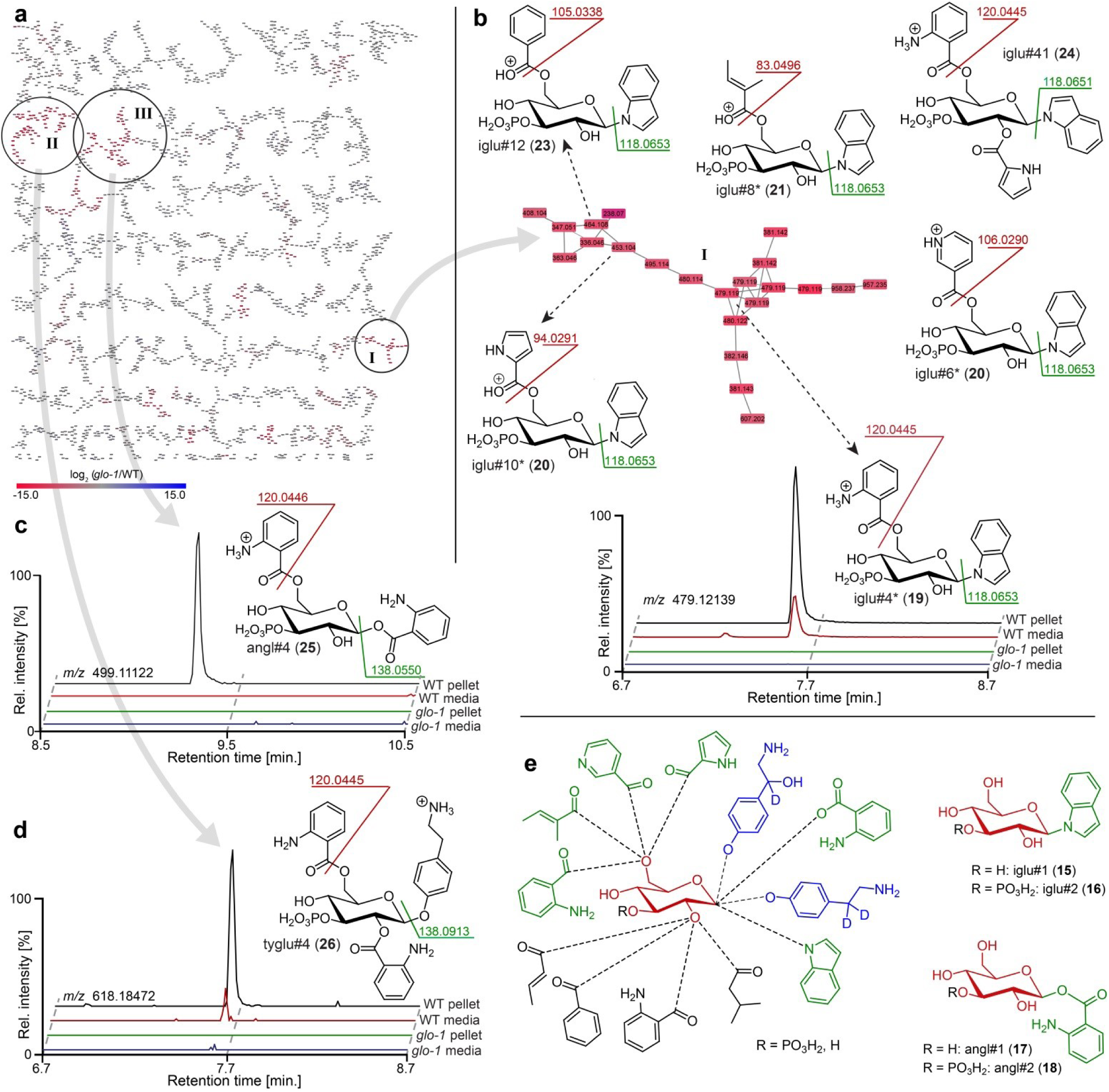
(a) Partial MS^2^ network (positive ion mode) for *C. elegans endo*-metabolome highlighting three clusters of modular glucosides that are down regulated in the *glo-1* mutants (also see Fig. S1-4). Red represents downregulated and blue upregulated features compared to wildtype *C. elegans*. (b) Cluster **I** feature several modular indole glucoside derivatives. Structures were proposed based on MS^2^ fragmentation patterns, also see Table 1. Compounds whose non-phosphorylated analogs were synthesized are marked (*). Shown ion chromatograms demonstrate loss of iglu#4 in *glo-1* mutants. (c,d) Examples for modular glucosides detected as part of clusters **II** and **III**. Ion chromatograms show abolishment of angl#4 (**25**) (c) and tyglu#4 (**26**) (d) production in *glo-1* mutants. (e) Modular glucosides are derived from combinatorial assembly of a wide range of building blocks. Incorporation of moieties was confirmed via total synthesis of example compounds (green) or stable isotope labeling (blue). For all compounds, 3-phosphorylation was proposed based on the established structures of iglu#2 (**16**), angl#2 (**18**), and uglas#11 (**3**).

As in the case of modular ascarosides, the abundances of putative building blocks of the newly identified modular glucosides were not strongly perturbed in *glo-1* mutants. For example, abundances of anthranilic acid, indole, octopamine, and tyramine were not significantly affected in *glo-1* null animals (Fig. S11). Notably, abundances of the glucosides scaffold, e.g. iglu#1 and angl#1, were also largely unaltered or even slightly increased in *glo-1* mutants (Fig. S11). In addition, production of some of the identified modular glucosides, e.g. iglu#5, is reduced but not fully abolished in *glo-1* worms (Fig. S8).

To confirm our results, we additionally compared the *glo-1* metabolome with that of *glo-4* mutants. *glo-4* encodes a predicted guanyl-nucleotide exchange factor acting upstream of *glo-1*, and like *glo-1* mutants, *glo-4* worms do not form LROs^18^. We found that the *glo-4* metabolome closely resembles that of *glo-1* worms, lacking most of the modular ascarosides and ascarosides detected in wildtype worms (Fig. S6c). Correspondingly, similar sets of compounds are upregulated in *glo-1* and *glo-4* mutants relative to wildtype, including ascarosyl glucosides and ascaroside phosphates. Compounds accumulating in *glo-1* and *glo-4* mutant worms further include a diverse array of small peptides (primarily three to six amino acids), consistent with the proposed role of LROs in the breakdown of peptides derived from proteolysis (Fig. S12)^36^. Taken together, our results indicate that, in addition to their roles in the degradation of metabolic waste, the LROs serve as hotspots of biosynthetic activity, where building blocks from diverse metabolic pathways are attached to glucoside and ascaroside scaffolds (Fig. 1a).

### Carboxylesterases are required for modular assembly

Comparing the relative abundances of different members of the identified families of modular glucosides and ascarosides, it appears that combinations of different building blocks and scaffolds are highly specific, suggesting the presence of dedicated biosynthetic pathways. For example, uric acid glucoside, gluric#1 (**6**), is preferentially combined with an ascaroside bearing a 7-carbon side chain (to form uglas#11, **3**), whereas ascarosides bearing a 9-carbon side chain are preferentially attached to the anomeric position of free glucose, as in glas#10 (**14**)^2,8^. Similarly, tiglic acid is preferentially attached to indole and tyramine glucosides but not to anthranilic acid glucosides (Table 1). Given that 4′- modification of ascarosides in *P. pacificus* and *C. elegans* require *cest* homologs, we hypothesized that the biosynthesis of other modular ascarosides as well as the newly identified glucosides may be under the control of *cest* family enzymes^26, 28^. From a list of 44 *uar-1* homologs from BLAST analysis (Table 2), we selected seven for further study (Fig. 3a, S2). The selected homologs are predicted to have intestinal expression, one primary site of small molecule biosynthesis in *C. elegans*^2^, and are closely related to the UAR-1 gene, while representing different sub-branches of the phylogenetic tree. Utilizing a recently optimized CRISPR/Cas9 method, we obtained two null mutant strains for five of the selected genes^37^. Mutants for the remaining two homologs, *ges-1* and *cest-6*, had been previously obtained (Table 3). We then analyzed the *exo*- and *endo*-metabolomes of this set of mutant strains by HPLC-HRMS to identify features that are absent or strongly downregulated in null mutants of a specific candidate gene compared to wildtype worms and all other mutants in this study. We found that two of the seven tested homologs (*cest-1.1*, *cest-2.2*) are defective in the production of two different families of modular ascarosides, whereas *cest-4* mutants were defective in the biosynthesis of a specific subset of modular indole glucosides (Fig. 3). The metabolomes of mutants for the remaining four *cest* homologs did not exhibit any significant differences compared to wildtype under the tested conditions.

**Figure 3:**
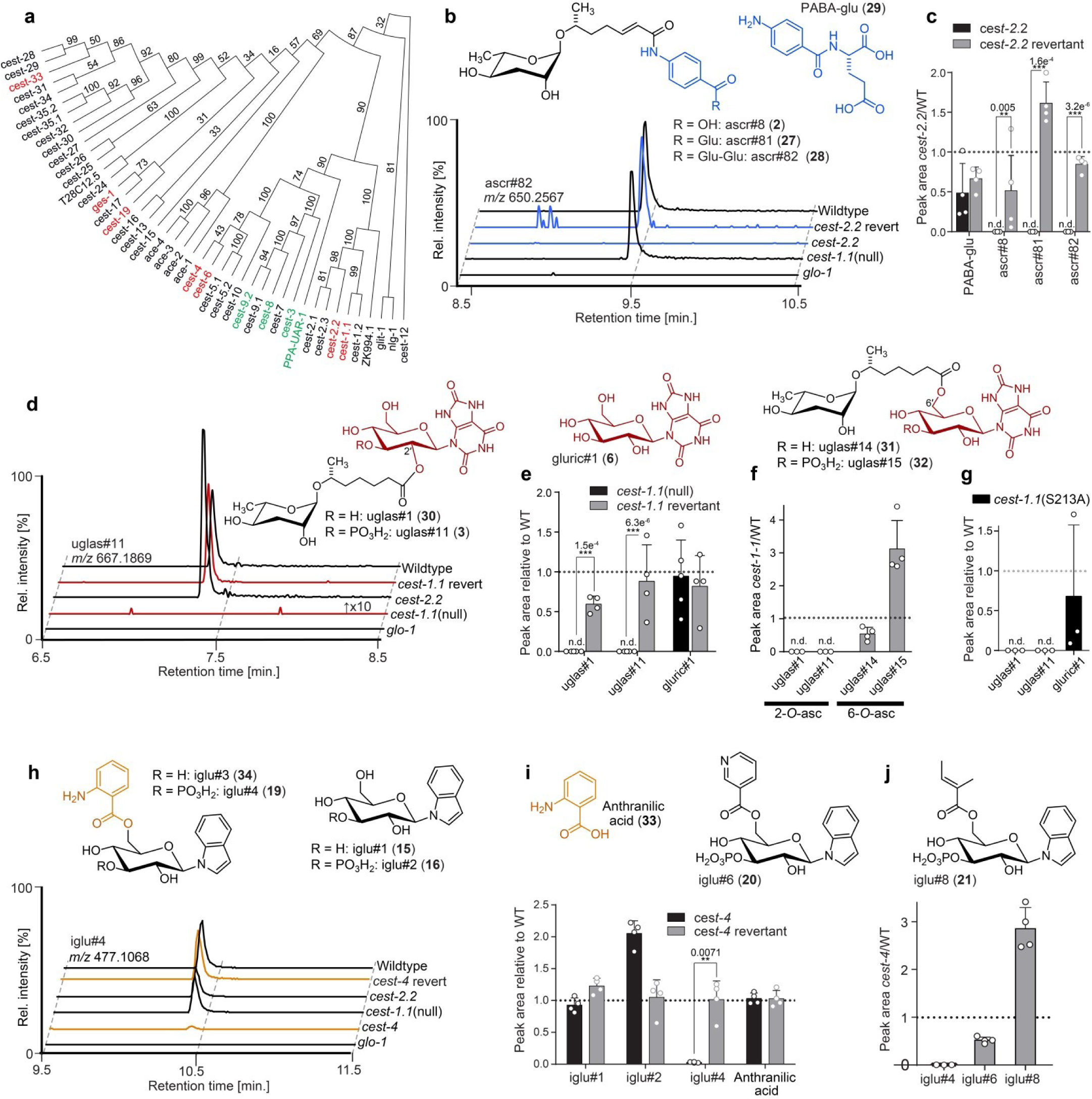
(a) Phylogenetic tree relating *P. pacificus uar-1* to homologous predicted genes in *C. elegans*. *Ppa-uar-1*, *cest-3*, *cest-8*, *cest-9.2* (green) mediate ester formation at the 4′-position of ascarosides in *P. pacificus* and *C. elegans*. Genes shown in red color were selected for the current study. (b,c) Production of ascr#8 (**2**), ascr#81 (**27**), and ascr#82 (**28**) is abolished in *cest-2.2* mutants Isogenic revertant strains of the *cest-2.2* null mutants in which the STOP-IN cassette was precisely excised, demonstrate wildtype-like recovery of the associated metabolite. (d,e) Production of uglas#1 and uglas#11 is abolished in *cest-1.1*(null) mutants and recovered in genetic revertants. (f) Biosynthesis of positional isomers uglas#14 (**31**) and uglas#15 (**32**) is unaltered or increased in *cest-1.1* mutants (f). (g) Production of uglas#1 and uglas#11, but not gluric#1, is abolished in *cest-1.1*(S213) mutants. (h,i) Production of the anthranilic acid-modified glucoside iglu#4 is largely abolished in *cest-4* mutants and fully recovered in genetic revertants. (j) Production of iglu#6 (**36**) and iglu#8 (**37**), whose structures are closely related to that of iglu#4, is not abolished in *cest-4* mutants. Ion chromatograms in panels b, d, and g further demonstrate abolishment in *glo-1* mutants. n.d., not detected. Error bars are standard deviation of the mean, and p-values are depicted in the Figure.

**Table 1.** MS^2^ data of glo-1 dependent features presented in this manuscript. Attached as a separate file.

**Table 2.** BLASTp results from the WormBase BLAST engine when searching against the amino acid sequence of UAR-1 and CRISPR/Cas9 targets for this study (red).

| Sequence | Score | E-value |
| --- | --- | --- |
| C01B10.10 | 280 | 2e-75 |
| C01B10.4a | 260 | 2e-69 |
| **T22D1.11** | **248** | **7e-66** |
| C42D4.2 | 233 | 4e-61 |
| **C17H12.4** | **231** | **1e-60** |
| C23H4.4a | 225 | 8e-59 |
| C23H4.7 | 199 | 6e-51 |
| C23H4.3 | 194 | 1e-49 |
| E01G6.3 | 193 | 3e-49 |
| C23H4.2 | 168 | 1e-41 |
| **T02B5.1** | **157** | **2e-38** |
| F15A8.6a | 154 | 1e-37 |
| F15A8.6b | 154 | 1e-37 |
| ZC376.3 | 153 | 3e-37 |
| T02B5.3 | 150 | 2e-36 |
| **ZC376.2b** | **148** | **1e-35** |
| **ZC376.2a** | **147** | **2e-35** |
| F56C11.6b | 141 | 1e-33 |
| F56C11.6a | 137 | 2e-32 |
| Y71H2AM.13 | 136 | 5e-32 |
| ZC376.1 | 135 | 1e-31 |
| R173.3 r | 129 | 6e-30 |
| T07H6.1a | 127 | 2e-29 |
| T28C12.4a | 124 | 1e-28 |
| T28C12.4b | 124 | 2e-28 |
| K07C11.4 | 119 | 6e-27 |
| **R12A1.4** | **118** | **1e-26** |
| K11G9.2 | 116 | 4e-26 |
| 02B12.4 | 115 | 8e-26 |
| Y75B8A.3 | 114 | 3e-25 |
| Y48B6A.8 | 113 | 4e-25 |
| F13H6.3 | 111 | 2e-24 |
| Y48B6A.7 | 109 | 5e-24 |
| 09B12.1 | 108 | 9e-24 |
| K11G9.1 | 108 | 2e-23 |
| ZC376.2c | 105 | 7e-23 |
| F07C4.12b | 105 | 7e-23 |
| C52A10.1 | 101 | 1e-21 |
| Y44E3A.2 | 101 | 2e-21 |
| K11G9.3 | 99 | 1e-20 |
| C52A10.2 | 97 | 3e-20 |
| C40C9.5d | 96 | 6e-20 |
| C40C9.5b | 96 | 6e-20 |
| C40C9.5a | 96 | 6e-20 |
| F55D10.3 | 96 | 1e-19 |
| C40C9.5f | 94 | 2e-19 |
| C01B10.4b | 94 | 2e-19 |
| C40C9.5g | 94 | 2e-19 |
| C40C9.5c | 94 | 3e-19 |
| C40C9.5e | 94 | 3e-19 |
| B0238.7 | 93 | 4e-19 |
| B0238.1 | 92 | 1e-18 |
| F55F3.2b | 83 | 6e-16 |
| F55F3.2a | 83 | 7e-16 |
| C23H4.4b | 50 | 5e-06 |
| Y43F8A.3a | 42 | 0.002 |
| Y43F8A.3b | 35 | 0.18 |

**Table 3.** List of *C. elegans* strains used in this study.

| Strain Name | Identifier | Description | Associated Metabolites |
| --- | --- | --- | --- |
| PS8031 | *cest-1.1 (sy1180)* | *cest-1.1* null | uglas#1 uglas#11 |
| PS8032 | *cest-1.1 (sy1181)* | *cest-1.1* null | uglas#1 uglas#11 |
| PS8259 | *cest-1.1 (sy1180 sy1250)* | *cest-1.1* null reverted to WT sequence | uglas#1 uglas#11 |
| PS8260 | *cest-1.1 (sy1180 sy1251)* | *cest-1.1* null reverted to WT sequence | uglas#1 uglas#11 |
| PS8261 | *cest-1.1 (sy1181 sy1252)* | *cest-1.1* null reverted to WT sequence | uglas#1 uglas#11 |
| PS8262 | *cest-1.1 (sy1181 sy1253)* | *cest-1.1* null reverted to WT sequence | uglas#1 uglas#11 |
| PS8008 | *cest-2.2 (sy1170)* | *cest-2.2* null | ascr#8, ascr#81, ascr#82 |
| PS8009 | *cest-2.2(sy1171)* | *cest-2.2* null | ascr#8, ascr#81, ascr#82 |
| PS8236 | *cest-2.2(sy1170 sy1236)* | *cest-2.2* null reverted to WT sequence | ascr#8, ascr#81, ascr#82 |
| PS8238 | *cest-2.2(sy1171 sy1238)* | *cest-2.2* null reverted to WT sequence | ascr#8, ascr#81, ascr#82 |
| PS8116 | *cest-4 (sy1192)* | *cest-4* null | iglu class modular glucosides |
| PS8117 | *cest-4(sy1193)* | *cest-4* null | iglu class modular glucosides |
| JJ1271 | *glo-1 (zu437)* | *glo-1* null | Most known modular ascarosides/glucosides |
| PS8781 | *cest-4 (sy1192)* | *cest-4* null reverted to WT sequence | iglu class modular glucosides |
| PS8782 | *cest-4 (sy1193)* | *cest-4* null reverted to WT sequence | iglu class modular glucosides |
| PS8783 | *cest-4 (sy1194)* | *cest-4* null reverted to WT sequence | iglu class modular glucosides |
| PS8784 | *cest-4 (sy1195)* | *cest-4* null reverted to WT sequence | iglu class modular glucosides |
| PS8515 | CBR-*glo-1*-A (sy1382) | *C. briggsae glo-1* null | Most known modular ascarosides/glucosides |
| PS8516 | CBR-*glo-1*-B (sy1383) | *C. briggsae glo-1* null | Most known modular ascarosides/glucosides |
| PS8029 | *cest-19(sy1178)* | *cest-19* null | Undetermined |
| PS8030 | *cest-19(sy1179)* | *cest-19* null | Undetermined |
| PS8033 | *cest-33(sy1182)* | *cest-33* null | Undetermined |
| PS8034 | *cest-33(sy1183)* | *cest-33* null | Undetermined |
| RB2053 | *ges-1 (ok2716)* | *ges-1* null | Undetermined |
| RB1804 | *cest-6(ok2338)* | *cest-6* null | Undetermined |
| DP683 | *cest-1.1(dp683)* | *cest-1.1* (S213A) point mutant | uglas#1 uglas#11 |
| FCS02 | *cest-2.2-mCherry* | *cest-2.2* C-terminal mCherry | ascr#8, ascr#81, ascr#82 |

**Table 4.**
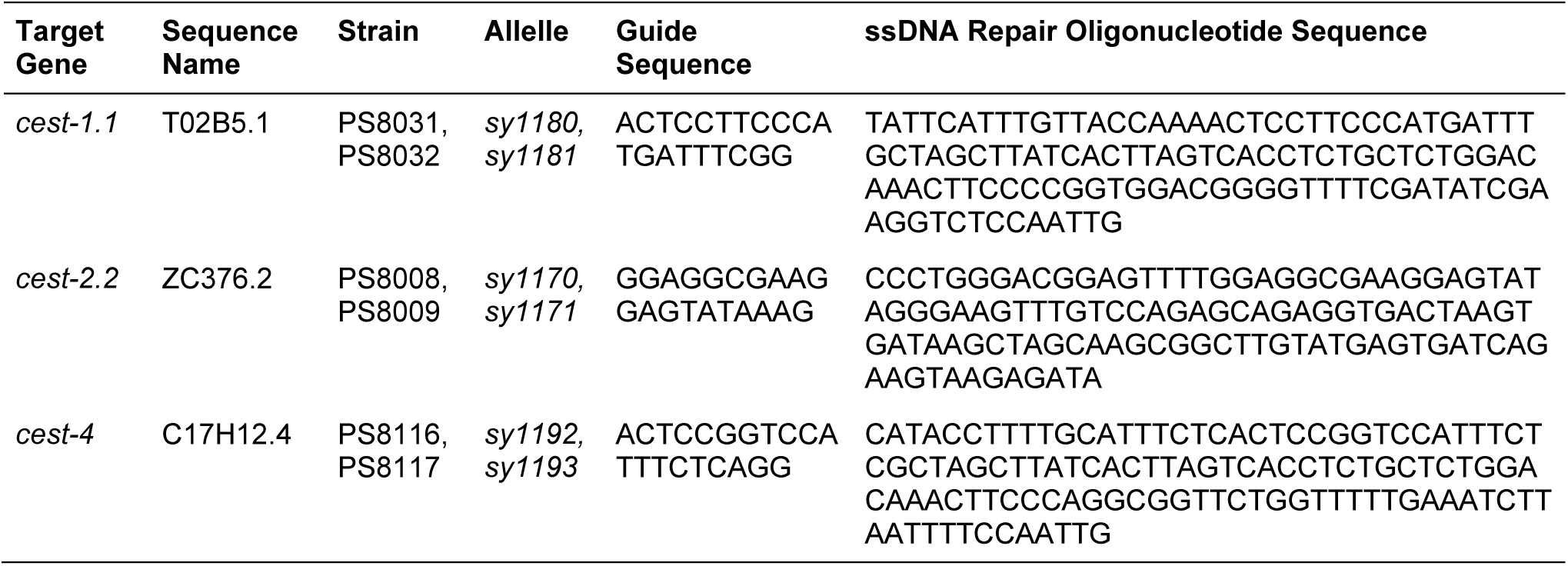
DNA oligonucleotides used for this study.

Analysis of the metabolomes of the two *cest-2.2* null mutants revealed loss of dauer pheromone component and male attractant ascr#8 (**2**) as well as of the closely related ascr#81 (**27**) and ascr#82 (**28**) (Fig. 3b, S13a). Biosynthetically, the ascr#8 family of ascarosides are derived from amide formation between ascr#7 (ΔC7, **7**) and folate-derived *p*-aminobenzoic acid (PABA, **8**), PABA-glutamate (**29**), or PABA-diglutamate, respectively. We did not detect any significant reduction in the production of plausible ascr#8 precursors, including PABA and PABA-glutamate, or ascr#7 (Fig. 3c, S14b). Biosynthesis of ascr#8, ascr#81, and ascr#82 was recovered in *cest-2.2* mutant worms in which the *cest-2.2* sequence had been restored to wild type using CRISPR/Cas9 (Fig. 3c, S15b). These results indicate that CEST-2.2 is required specifically for biosynthesis of the amide linkage between the carboxy terminus of ascr#7 and PABA derivatives, in contrast to the implied functions of UAR-1, CEST-8, CEST-3, and CEST- 9.2, which are involved in the formation of ester bonds between various head groups and the 4′- hydroxy group of ascarylose^26, 28^.

In *cest-1.1* null mutants (*cest-1.1*(null)), biosynthesis of the nucleoside-like ascaroside uglas#1 (**30**) and its phosphorylated derivative uglas#11 (**3**) was abolished (Fig. 3d, S13c). uglas#1 and uglas#11 are derived from the attachment of ascr#1, bearing a seven carbon (C7) side chain, to the uric acid gluconucleoside gluric#1 (**6**). Production of ascr#1 (**5**) and gluric#1 (**6**), representing plausible building blocks of uglas#1 (**30**), was not reduced (Fig. S14a). Furthermore, production of uglas#14 (**31**) and uglas#15 (**32**), isomers of uglas#1 and uglas#11 bearing the ascarosyl moiety at the 6′ position instead of the 2′ position, was not abolished but rather slightly increased in *cest-1.1*(null) (Fig. 3d-e). These results indicate that CEST-1.1 is required for the formation of the ester bond specifically between ascr#1 (**5**) and the 2′-hydroxyl group in gluric#1. As in the case of *cest-2.2*, biosynthesis of uglas#1 and uglas#11 was fully recovered in *cest-1.1* mutant worms in which the *cest-1.1* sequence had been restored to wild type using CRISPR/Cas9 (Fig. 3f, S15a).

Sequence alignment with human AChE suggested that serine 213 is part of the conserved catalytic serine-histidine-glutamate triad of CEST-1.1 (Fig. S16). To test whether disruption of the catalytic triad would affect production of *cest-1.1-*dependent metabolites, we generated a point mutant, *cest-1.1*(S213A). As in *cest-1.1*(null), production of uglas#1 (**30**) and uglas#11 (**3**) was fully abolished in *cest-1.1*(S213A), whereas production of gluric#1 was not affected (Fig. 3g).

Previous work implicated *cest-1.1* with longevity phenotypes associated with argonaute-like gene 2 (*alg-2*)^38^. *alg-2* mutant worms are long lived compared to wild type and their long lifespan was further shown to require *daf-16*, the sole ortholog of the FOXO family of transcription factors in *C. elegans*, as well as *cest-1.1*. Moreover, uglas#11 biosynthesis is significantly increased in mutants of the insulin receptor homolog *daf-2*, a central regulator of lifespan in *C. elegans* upstream of *daf-16*^12^. These findings suggest the possibility that the production of uglas ascarosides underlies the *cest-1.1*-dependent extension of adult lifespan in *C. elegans*.

In contrast to our results for *cest-1.1* and *cest-2.2* mutants, comparative metabolomic analysis of the *cest-4* mutant strains did not reveal any defects in the biosynthesis of known ascarosides. Instead, we found that the levels of a specific subset of modular anthranilic acid (**33**) bearing indole glucosides, including iglu#3 (**34**) and its phosphorylated derivative iglu#4 (**35**) were abolished in the *cest-4* mutant worms (Fig. 3h, S13b). Abundances of the putative precursor glucosides, iglu#1 (**15**) and iglu#2 (**16**), were not significantly changed in *cest-4* (Fig. 3i, S14c). Notably, production of other indole glucosides, e.g. iglu#6 (**36**) and iglu#8 (**37**), was not significantly reduced in *cest-4* worms (Fig. 3i, S17). Biosynthesis of iglu#3 and iglu#4 was restored to wild type levels in genetic revertant strains for *cest-4* (Fig. 3i, S15c). Therefore, it appears that *cest-4* is specifically required for attachment of anthranilic acid to the 6′ position of glucosyl indole precursors, whereas attachment of tiglic acid, nicotinic acid, and other moieties is *cest-4*-independent (Fig. 3j, S17). The role of *cest-4* in the biosynthesis of the iglu family of modular glucosides thus parallels that of *cest-1.1* in the biosynthesis of the uglas ascarosides: whereas *cest-4* appears to be required for the attachment of anthranilic acid (**33**) to the 6’ position of a range of indole glucosides, *cest-1.1* appears to be required for attaching the ascr#1 side chain to the 2′ position in uric acid glucosides.

### CEST-2.2 localizes to intestinal granules

All *cest* homologs selected for this study exhibit domain architectures typical of the α/ß-hydrolase superfamily of proteins, including a conserved catalytic triad, and further contain a predicted disulfide bridge, as in mammalian AChE^39^ (Fig. S16). The *cest* genes also share homology with neuroligin, a membrane bound member of the α/ß-hydrolase fold family, that mediates the formation and maintenance of synapses between neurons^40^. Sequence analysis suggests that five of the seven CEST homologs studied here are membrane anchored (Fig. S18), given the presence of a predicted *C*-terminal transmembrane domain^41^ (consisting of ∼20 residues), with the *N* terminus on the luminal side of a vesicle or organelle (Fig. S18). Since the production of all so far identified *cest*-dependent metabolites is abolished in *glo-1* mutants, it seemed likely that the CEST proteins localize to intestinal granules. To test this idea, we created a mutant strain that express *cest-2.2 C*-terminally tagged with mCherry at the native genomic locus to avoid potentially confounding effects of overexpression. The red fluorescent mCherry was chosen because of the strong green autofluorescence of the LROs^17^. We confirmed that production of all *cest-2.2*-dependent metabolites, including ascr#8 (**2**), ascr#81 (**27**), and ascr#82 (**28**) was not significantly altered in *cest-2.2*-mCherry mutants (Fig. 4a), indicating that CEST-2.2 remained functional. Imaging of wildtype adult worms revealed strong green and weaker red autofluorescence in circular features in intestinal cells, consistent with LROs. In addition, *cest-2.2*-mCherry-tagged worms showed red fluorescence in a distinct set of intestinal granules that showed little if any autofluorescence (Fig. 4b, S19, S20). It is unclear whether mCherry also localizes to the strongly autofluorescent granules, as we cannot distinguish the mCherry signal from the red component of the autofluorescence, given relatively low CEST-2.2-mCherry expression in this non-overexpressing strain. Taken together, it appears that CEST-2.2-mCherry localizes to a subset of intestinal organelles that is partly distinct from the autofluorescent LROs. Further studies are required to determine if CEST-2.2-mCherry co-localizes with other intestinal granule markers, specifically GLO-1 and the lysosomal marker LMP-1.

**Figure 4:**
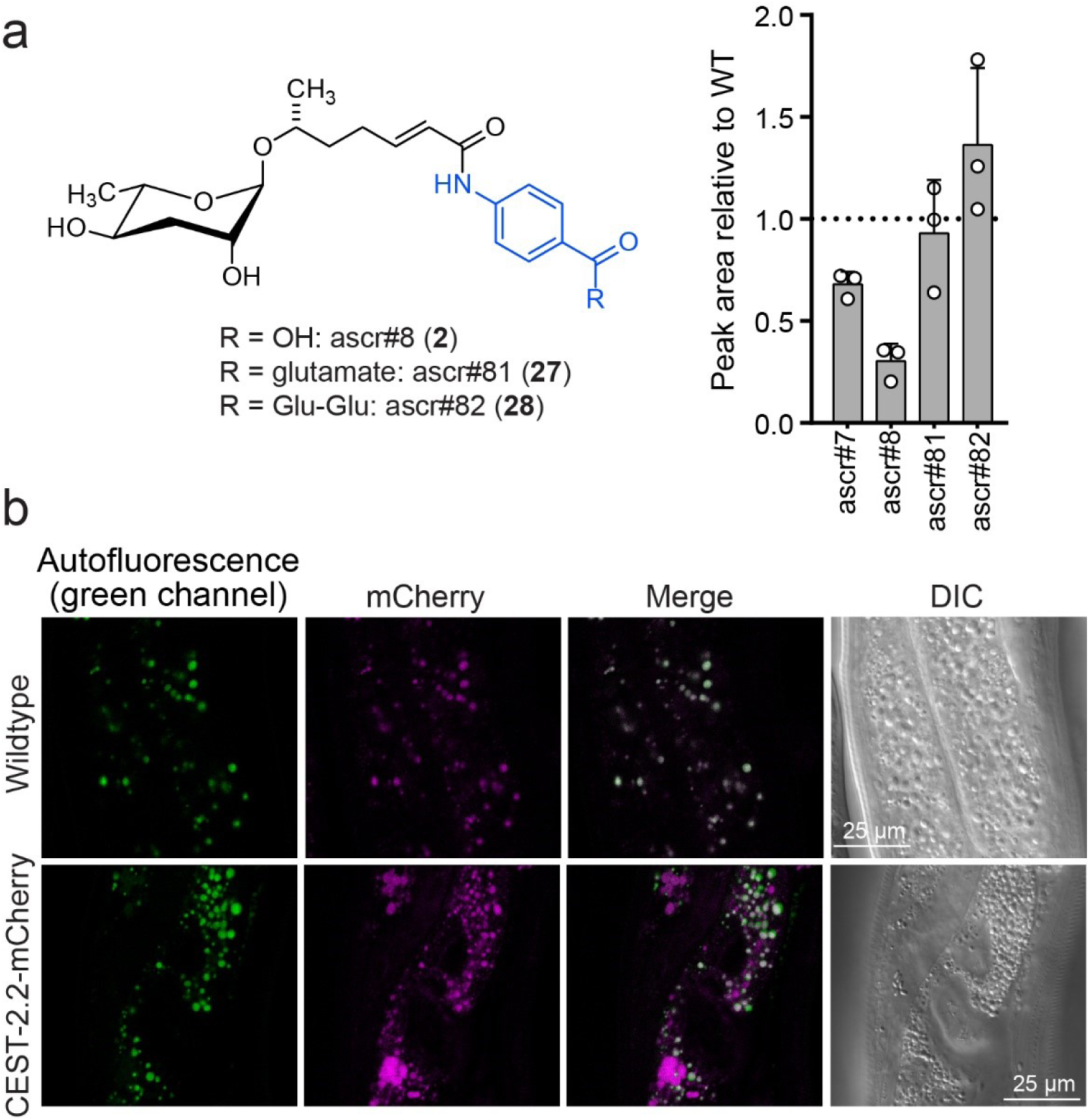
(a) Relative amounts of *cest-2.2* dependent metabolites in worms expressing *C*- terminally mCherry-tagged CEST-2.2. (b) Red fluorescence in intestinal granules in wildtype and *cest-2.2*-mCherry gravid adults. Top, wildtype (N2) control; bottom, *cest-2.2*-mCherry worms.

### *Glo-1*-dependent metabolites in *C. briggsae*

In addition to *C. elegans* and *P. pacificus*, modular ascarosides have been reported from several other *Caenorhabditis* species^42, 43^, including *C. briggsae*^23, 44^. To assess whether the role of LROs in the biosynthesis of modular metabolites is conserved across species, we created two *Cbr*-*glo-1* (CBG01912.1) knock-out strains using CRISPR/Cas9. As in *C. elegans*, *Cbr*-*glo-1* mutant worms lacked autofluorescent LROs, which are prominently visible in wildtype *C. briggsae* (Fig. S21). Comparative metabolomic analysis of the *endo*- and *exo*-metabolomes of wildtype *C. briggsae* and the *Cbr*- *glo-1* mutant strains revealed that biosynthesis of all known modular ascarosides is abolished in *Cbr*-*glo-1* worms, including the indole carboxy derivatives icas#2 (**35**) and icas#6.2 (**36**), which are highly abundant in wildtype *C. briggsae* (Fig. 5a).^23^ In addition, the *C. briggsae* MS^2^ networks included several large *Cbr*-*glo-1*-dependent clusters representing modular glucosides, including many of the compounds also detected in *C. elegans*, e.g. iglu#4 and angl#4. As in *C. elegans*, production of unmodified glucoside scaffolds, e.g. iglu#1 (**15**) and angl#1 (**17**), was not reduced or increased in *Cbr*-*glo-1* mutants, whereas biosynthesis of most modular glucosides derived from attachment of additional moieties to these scaffolds was abolished (Fig. 5b). Taken together, these results indicate that the role of LROs as a central hub for the assembly of diverse small molecule architectures, including modular glucosides and ascarosides, may be widely conserved among nematodes (Fig. 5c).

**Figure 5:**
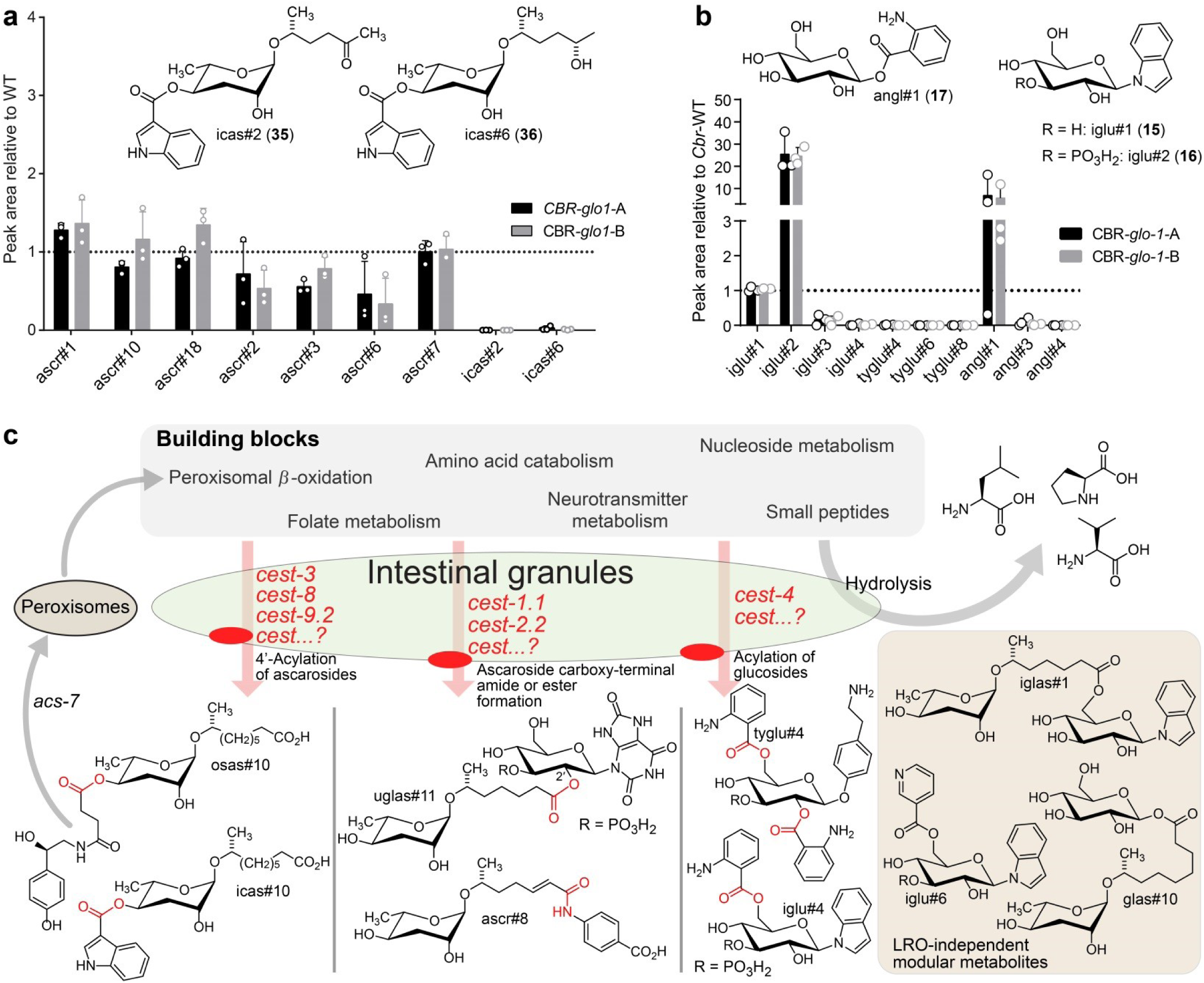
Relative abundance of (a) simple and modular ascarosides and (b) simple and modular glucosides in the *endo*-metabolome of *Cbr*-*glo-1* mutants relative to wildtype *C. briggsae*. n.d., not detected. (c) Model for modular metabolite assembly. CEST proteins (membrane-bound in the LROs, red) mediate attachment of building blocks from diverse metabolic pathways to glucose scaffolds and peroxisomal *β*-oxidation-derived ascarosides via ester and amide bonds. Some of the resulting modular ascarosides may undergo additional peroxisomal *β*-oxidation following activation by *acs-7*^25^.

## Discussion

Our results indicate that in *C. elegans* the Rab-GTPase *glo-1,* which is required for formation of intestinal LROs, plays a central role in the biosynthesis of several large compound families derived from modular assembly via *cest* homologs. Formation of the autofluorescent LROs via *glo-1* is reminiscent of the roles of its human orthologs RAB32 and RAB38, which are required for the formation of melanosomes, and perhaps other LROs^45, 46^. Lysosomes and LROs are generally presumed to function in autophagy, phagocytosis, and the hydrolytic degradation of proteins, and Rab32 family GTPases have been shown to be required for these processes in diverse organisms^47^. Consistent with the notion that lysosomes and LROs are degradation hotspots, many of the building blocks of the identified modular ascarosides and glucosides are derived from catabolic pathways, for example, anthranilic acid is derived from tryptophan catabolism, uric acid stems from purine metabolism, and the short chain ascarosides are the end products of peroxisomal *β*-oxidation of very long-chain precursors. Importantly, although our results indicate that carboxylesterases participate in *glo-1*-dependent modular metabolite assembly, additional studies are required to clarify whether the intestinal compartments that carboxylesterases localize to also contain GLO-1 and the lysosomal marker LMP-1, as is the case for the autofluorescent LROs^22^.

Further, our results demonstrate that the modular assembly paradigm extends beyond ascarosides. The modular glucosides represent a previously unknown family of nematode metabolites. In contrast to the well-established role of modular ascarosides as pheromones, it is unknown whether modular glycosides serve specific biological functions, e.g., as signaling molecules; however, their specific biosynthesis via *cest-4* as well as their life stage-dependent production strongly supports this hypothesis (Fig. S10). Like the ascaroside pheromones, some modular glucosides are excreted into the media, suggesting that they could be involved in inter- organismal communication. Identifying developmental and environmental conditions that affect modular glucoside production, as well as a more comprehensive understanding of their biosyntheses, may help uncover potential signaling and other biological roles. In particular, the apparent peroxisomal origin of the ascaroside scaffolds suggests a link between peroxisome and gut granule activity, perhaps via pexophagy^48^, and characterization of the role of autophagy for gut granule-dependent metabolism may contribute to uncovering the functions of modular glucoside and ascarosides. A connection to autophagy is also suggested by our previous finding^14^ that production of modular ascarosides is reduced in mutants of *atg-18*^49^, which is essential for autophagy.

The high degree of selectivity in which different building blocks are combined in the modular ascarosides and glucosides strongly suggests that these compounds, despite their numbers and diversity, represent products of dedicated enzymatic pathways, as has recently been established for 4′-acylated ascarosides. Our results revealed a wider range of biosynthetic functions associated with *cest* homologs, including esterification and amide formation at the carboxy terminus of ascarosides and acylation of glucosides (Figure 5c). Notably, all *cest* null mutants whose metabolomes have been characterized so far are defective in the biosynthesis of one or a few compounds sharing a specific structural feature, further supporting the view that these selectively assembled molecular architectures serve dedicated functions.

All CEST proteins that so far have been associated with modular metabolite assembly contain membrane-anchors and exhibit domain architectures typical of serine hydrolases of the AChE family, including an α/β-hydrolase fold, a conserved catalytic serine-histidine-glutamate triad, and bridging disulfide cysteines (Fig. S16)^39^. While our efforts at heterologous expression of CEST proteins were unsuccessful, the finding that mutation of the catalytic serine in *cest- 1.1*(S213A) abolished production of all *cest-1.1*-dependent compounds suggests that CEST enzymes directly participate in the biosynthesis of modular metabolites. Therefore, we hypothesize that CEST proteins, after translating from the endomembrane system to *glo-1*- dependent intestinal organelles, partake in the assembly of diverse ascaroside or glucoside- based architectures via acyl transfer from corresponding activated intermediates, e.g. CoA or phosphate esters^39, 50^. α/β-hydrolase fold enzymes are functionally highly diverse^51^ and include esterases, peptidases, oxidoreductases, and lyases, serving diverse biosynthetic roles in animals, plants^52^, and bacteria^53^. While acyltransferase activity is often observed as a side reaction for esterases and lipases, α/β-hydrolase fold enzymes can function as dedicated acyltransferases, e.g. in microbial natural product biosyntheses^51, 54^. Additional biochemical studies will be required to delineate the exact mechanisms by which *cest* homologs contribute to modular metabolite assembly in nematodes.

Finally, although our results indicate that *glo-1* is required for the biosynthesis of most modular metabolites we have detected so far, it is notable that some modular ascarosides, e.g. iglas#1 (**13**), and modular glucosides, e.g. iglu#6 (**20**) and iglu#8 (**21**), do not appear to be *glo- 1*-dependent (Fig. S8). This suggests that diverse cell compartments contribute to modular metabolite biosynthesis and may also indicate that not all CEST proteins are delivered to the same cellular compartment. Similarly, *glo-1* mutants continue to generate the simple glucosides and ascarosides that serve as scaffolds for further elaboration via CEST proteins, which may be derived from UDP-glycosyltransferases^55^.

Reminiscent of the role of AChE for neuronal signal transduction in animals, it appears that, in *C. elegans*, carboxylesterases with homology to AChE have been co-opted to establish additional signal transduction pathways that are based on a modular chemical language, for inter-organismal communication, and perhaps also intra-organismal signaling. The biosynthetic functions of most of the 200 serine hydrolases in *C. elegans*, including more than 30 additional *cest* homologs, remain to be assessed, and it seems likely that this enzyme family contributes to the biosynthesis of a large number of additional, yet unidentified compounds. Similarly, the exact enzymatic roles of many families of mammalian serine hydrolases have not been investigated using HRMS-based untargeted metabolomics. Our results may motivate a systematic characterization of metazoan *cest* homologs and other serine hydrolases, with regard to their roles in metabolism and small molecule signaling, associated enzymatic mechanisms, and cellular localization.

## Methods

### General information

Unless noted otherwise, all reagents were purchased from Sigma-Aldrich. All newly identified compounds were assigned four letter “SMID”s (a search-compatible, Small Molecule IDentifier) e.g., “icas#3” or “ascr#10”. The SMID database (www.smid-db.org) is an electronic resource maintained in collaboration with WormBase (www.wormbase.org). A complete list of SMIDs can be found at www.smid-db.org/browse, and example structures for different SMIDs at www.smid-db.org/smidclasses.

### BLAST analysis of *uar-1*

Amino acid sequence of *Ppa*-UAR-1 was used as previously published^26^. BLASTp was run from the WormBase engine at (https://wormbase.org/tools/blast_blat). E-value threshold was set to 1E0. Database was set to WS269 and species was set to *C. elegans*. Results of BLASTp search are listed in Table 2.

### Amino acid sequence alignment

hAChE was aligned with *Ppa*-UAR-1, CEST-1.1, CEST-2.2, and CEST-4 was done using T-Coffee Multiple Sequence alignment^56^. Protein sequences for *C. elegans* CEST proteins are from WormBase. The AChE sequence was obtained from NCBI (accession number P22303). Amino acids were colored based on chemical properties: AVFPMILW = red (small + hydrophobic), DE = blue (acidic), RHK = magenta (basic), STYHCNGQ = green (hydroxyl + sulfhydryl + amine + glycine). See Figure S17 for results.

### Phylogenetic tree

The protein sequence of Ppa-UAR1 was submitted to an NCBI BLASTp search^57^ (restricted to species *C. elegans*, conditional compositional BLOSUM62, gap open cost:11, gap extension cost: 1, word size: 6) using Geneious software (Biomatters Inc). The top BLAST hits by E-value up to and including *ace-3* were selected, and only the best scoring transcript variant was kept for each protein sequence hit. A total of 28 sequences were then imported into MEGA7^58^ and aligned using MUSCLE^59^ (settings: gap open penalty: −2.9, gap extend 0, hydrophobicity multiplier 1.2, max. iterations 8, clustering method for all iterations: UPGMB, minimal diagonal length: 24). From this alignment, an Maximum Likelihood tree was built based on the JTT matrix-based model^60^. Initial trees were built by applying Neighbor-Join and BioNJ algorithms to a matrix of pairwise distances estimated using a JTT model assuming uniform substitution rates across positions. Phylogeny confidence was tested using 200 bootstrap replications. The tree with the highest log likelihood (−22299.9282) is shown. At each branch, the percentage of bootstrap replicates containing the same branching event is denoted. The tree is drawn to scale, with branch lengths measured in the number of substitutions per site. The evolutionary history was inferred by using the Maximum Likelihood method based on the JTT matrix-based model^60^. The tree with the highest log likelihood (−22299.9282) is shown. The percentage of trees in which the associated taxa clustered together is shown next to the branches. Initial tree(s) for the heuristic search were obtained automatically by applying Neighbor-Join and BioNJ algorithms to a matrix of pairwise distances estimated using a JTT model, and then selecting the topology with superior log likelihood value. The tree is drawn to scale, with branch lengths measured in the number of substitutions per site. The analysis involved 28 amino acid sequences. All positions containing gaps and missing data were eliminated. There were a total of 427 positions in the final dataset. Evolutionary analyses were conducted in MEGA7^58, 61^.

### Nematode strains

Wildtype (N2) and *glo-1(zu437)* null animals were provided by the Caenorhabditis Genetics Center (CGC), which is funded by NIH Office of Research Infrastructure Programs (P40 OD010440). *cest-2.2* mutant strain integrating N-terminal (mCherry-*cest-2.2*) or C-terminal mCherry (*cest-2.2*-mCherry), were generated by SunyBiotech. Generation of *C. elegans and C. briggsae* null mutants and revertants as well as generation of the *cest-1.1* point mutant is described below. See Table 3 for a complete list of strains used in this study.

### *C. elegans* CRISPR mutagenesis for generation of *cest* null mutants

CRISPR/Cas9 mutagenesis was performed as in *Wang et al., 2018*^37^. Briefly, *C. elegans* strain N2 were gene-edited by insertion of a 43-base-pair insertion that disrupts translation. Independent homozygous mutants were picked among the progeny of heterozygous F1 progeny of injected hermaphrodites and given distinct unique allele names. Reversion of mutants was accomplished in the same way.

### *C. briggsae* CRISPR mutagenesis for generation of *glo-1* null mutants

The *C. briggsae glo-1* mutants sy1382 and sy1383 were both created using the briggsae adaptation of the STOP-IN cassette method as described in *Cohen and Sternberg 2019*^62^ and *Wang et al. 2018*^37^. Both strains were made using a successful insertion of the STOP-IN cassette into the middle of the first exon using the guide AACAAATCTCCGGATGATTG. To detect the insertion, we used forward primer GGGTGACCGCCCATTTATTG and reverse primer AAAGGCGCACATCTTGCTTC.

### *C. elegans* CRISPR mutagenesis for generation of the *cest-1.1(dp683)* allele encoding the S213A catalytic mutant

*cest-1(dp683)* was generated as previously described^63^. Briefly, *daf- 2(e1368)* mutant animals were injected with *in-vitro-*assembled Cas9-crRNA-tracrRNA complexes targeting *cest-1.1* and the *dpy-10* co-CRISPR gene and two 100bp repair oligonucleotides containing the desired *cest-1.1* mutation and the *dpy-10(cn64)* co-CRISPR mutation^64^. Sequences of the *cest-1.1* crRNA and repair oligonucleotide are 5’ acctacCGCTACTATCATAC 3’ and 5’ GAAATTGAAAACTTTGGAGGAAATAAAAACAG- AATTACATTGGCAGGGCATGCCGCTGGAGCAAGTATGATAGTAGCGgtaggtcacataaatgataca tttttg 3’, respectively. F1 Rol progeny of injected animals were picked and screened for the presence of the *cest-1.1(dp683)* mutation after egglay. F2 broods of F1 Rol animals that were heterozygous for *cest-1.1(dp683)* were screened for animals that were homozygous for *cest- 1.1(dp683)* and either wild-type or heterozygous for *cn64* at the *dpy-10* locus. Subsequent broods were screened for wild-type *dpy-10* animals to remove the co-CRISPR mutation.

### Nematode imaging

To image, gravid adult *C. elegans* were transferred to an agarose pad on a glass slide with 10 µM of levamisole to immobilize the worms. Microscopic analysis was performed using a Leica TCS SP5 Laser Scanning Confocal Microscope. Green autofluorescence was excited at 488 nm and the emission detector was set to 490-540 nm. mCherry was excited with 561 nm and the emission detector was set to 590-650 nm. Worms were imaged using the 100x objective.

### C. briggsae imaging

0.5 mL of 2 µM Lysotracker Deep Red (Thermo Fisher 1 mM stock in DMSO) was added to a 6 cm NGM plate seeded with 0.1 mL of *E. coli* OP50 and incubated in the dark for 24 hours at 20 °C. L4 larvae of *C. briggsae* were added to the plate and allowed to grow in the dark for 24 hours at 20 °C. To image, *C. briggsae* were transferred to an agarose pad on a glass slide with 10 µM of levamisole to immobilize the worms. Microscopic analysis was performed using a Zeiss Axio Imager Z2 florescence microscope with Apotome.

### Nematode cultures, mixed stage

Culturing began by chunking *C. elegans* or *C. briggsae* onto 10 cm NGM plates (each seeded with 800 µL of OP50 *E. coli* grown to stationary phase in Lennox Broth) and incubated at 22 °C. Once the food was consumed, the cultures were incubated for an additional 24 hours. Each plate was then washed with 25 mL of S-complete medium into a 125 mL Erlenmeyer flask, and 1 mL of OP50 *E. coli* was added (*E. coli* cultures were grown to stationary phase in Terrific Broth, pelleted and resuspended at 1 g wet mass per 1 mL M9 buffer), shaking at 220 RPM and 22 °C. After 70 hours, cultures were centrifuged at 5000 G for 1 min. After discarding supernatant, 24 mL H_2_O was added, along with 6 mL bleach, 900 µL 10 M NaOH and the mixture was shaken for 3 min to prepare eggs. Eggs were centrifuged at 5000 G, the supernatant was removed, and the egg pellet washed with 35 mL M9 buffer twice and then suspended in a final volume of 5 mL M9 buffer in a 50 mL centrifuge tube. Eggs were counted and placed on a rocker and allowed to hatch as L1 larvae for 24 hours at 22 °C. 70,000 L1 larvae were seeded in 25 mL cultures of S-complete with 1 mL of OP50 and incubated at 220 RPM and 22 °C in a 125 mL Erlenmeyer flask. After 72 hours, cultures were fed an additional 1 mL of OP50 and incubation continued. After an additional 48 hours, worms were spun at 1000 G 5 min and spent medium was separated from worm body pellet. Separated medium and worm pellet were flash frozen over liquid nitrogen until further processing. At least three biological replicates were grown for all mutant strains. Mutants were grown with parallel wildtype controls, and biological replicates were started on different days.

### Metabolite extraction

Lyophilized pellet and media samples were crushed and homogenized by shaking with 2.5 mm steel balls at 1300 rpm for 3 min in 30 s pulses while chilled with liquid nitrogen (SPEX sample prep miniG 1600). Thus powdered media and pellet samples were extracted with 15 mL methanol in 50 mL centrifuge tubes, rocking overnight at 22 °C. Extractions were pelleted at 5000 g for 10 min at 4 °C, and supernatants were transferred to 20 mL glass scintillation vials. Samples were then dried in a SpeedVac (Thermo Fisher Scientific) vacuum concentrator. Dried materials were resuspended in 1 mL methanol and vortexed for 1 min. Samples were pelleted at 5000 g for 5 min and 22 °C, and supernatants were transferred to 2 mL HPLC vials and dried in a SpeedVac vacuum concentrator. Samples were then resuspended in 200 μL of methanol, transferred into 1.7 mL Eppendorf tubes, and centrifuged at 18,000 G for 20 min at 4 °C. Clarified extracts were transferred to fresh HPLC vials and stored at −20 °C until analysis.

### Preparation of *exo*-metabolome samples from staged starved and fed cultures

40,000 synchronized L1 larvae were added to 125 mL Erlenmeyer flasks containing 30 mL of S-complete medium. Worms were fed with 4 mL of concentrated OP-50 and incubated at 20 °C with shaking at 160 RPM for: 12 h (L1), 24 h (L2), 32 h (L3), 40 h (L4) and 58 h (gravid adults). For preparation of starved samples, each of the stages was starved for 24 h after reaching their desired developmental stage in S-complete without OP-50. After incubation for the desired time, liquid cultures were centrifuged (1000 x g, 22 °C, 1 min) and supernatants were collected. Supernatant was separated from intact OP-50 cells by centrifuging (3000 x g, 22 °C, 5 min) and the resulting supernatants (*exo*-metabolome) were lyophilized. Lyophilized samples were homogenized with a dounce homogenizer in 10 mL methanol and extracted on a stirring plate (22 °C, 12 h). The resulting suspension was centrifuged (4000 g, 22 °C, 5 min) to remove any precipitate before carefully transferring to an LC-MS sample vial. Three biological replicates were started on different days.

### Mass spectrometric analysis

High resolution LC-MS analysis was performed on a Thermo Fisher Scientific Vanquish Horizon UHPLC System coupled with a Thermo Q Exactive HF hybrid quadrupole-orbitrap high-resolution mass spectrometer quipped with a HESI ion source.

1 μL of extract was injected and separated using at water-acetonitrile gradient on a Thermo Scientific Hypersil GOLD C18 column (150 mm x 2.1 mm 1.9 um particle size 175 Å pore size, Thermo Scientific) and maintained at 40 °C. Solvents were all purchased from Fisher Scientific as HPLC grade. Solvent A: 0.1% formic acid in water; solvent B: 0.1% formic acid in acetonitrile. A/B gradient started at 1% B for 5 min, then from 1% to 100% B over 20 min, 100% for 5 min, then down to 1% B for 3 min. Mass spectrometer parameters: 3.5 kV spray voltage, 380 °C capillary temperature, 300 °C probe heater temperature, 60 sheath flow rate, 20 auxiliary flow rate, 1 spare gas; S-lens RF level 50.0, resolution 240,000, *m/z* range 100-1200 m/z, AGC target 3e6. Instrument was calibrated with positive and negative ion calibration solutions (Thermo-Fisher) Pierce LTQ Velos ESI pos/neg calibration solutions.

### Feature detection and characterization

LC−MS RAW files from each sample were converted to mzXML (centroid mode) using MSConvert (ProteoWizard), followed by analysis using the XCMS^65^ analysis feature in METABOseek (metaboseek.com). Peak detection was carried out with the centWave algorithm^29^, values set as: 4 ppm, 320 peakwidth, 3 snthresh, 3100 prefilter, FALSE fitgauss, 1 integrate, TRUE firstBaselineCheck, 0 noise, wMean mzCenterFun, -0.005 mzdiff. XCMS feature grouping values were set as: 0.2 minfrac, 2 bw, 0.002 mzwid, 500 max, 1 minsamp, FALSE usegroup. METABOseek peak filling values set as: 5 ppm_m, 5 rtw, TRUE rtrange. Resulting tables were then processed with the METABOseek Data Explorer. Molecular features were filtered for each particular null mutant against all other mutants. Filter values were set as: 10 to max minFoldOverCtrl, 15000 to max meanInt, 120 to 1500 rt, 0.95 to max Peak Quality as calculated by METABOseek. Features were then manually curated by removing isotopic and adducted redundancies. Remaining masses were put on the inclusion list for MS/MS (ddMS2) characterization. Positive and negative mode data were processed separately. In both cases we checked if a feature had a corresponding peak in the opposite ionization mode, since fragmentation spectra in different modes often provide complementary structural information. To acquire MS2 spectra, we ran a top-10 data dependent MS2 method on a Thermo QExactive-HF mass spectrometer with MS1 resolution 60000, AGC target 1 × 10^6, maximum IT (injection time) 50 ms, MS2 resolution 45 000, AGC target 5 × 10^5, maximum IT 80 ms, isolation window 1.0 m/z, stepped NCE (normalized collision energy) 25, 50, dynamic exclusion 3 s.

### Statistical analysis

Peak integration data from HPLC-MS analysis were log-transformed^66^ prior to statistical analysis. Significance of differences between average peak areas were then assessed using unpaired t-tests.

### MS^2^-based molecular networking

From the list of differential features, described above, MS^2^ data was acquired for these features. To generate the MS^2^ molecular network, Metaboseek version 0.9.6 was used. Using the MS2scans features, differential features were matched with their respective MS^2^ scan, using *m/z* window of 5 ppm, and a retention time window of 15 sec. To construct the molecular network, tolerance of the fragment peaks was set to *m/z* of 0.002 or 5 ppm, minimum number of peaks was set to 5, with a 2% noise level. Once the network was constructed, cos value of 0.8 was used, and the number of possible connections was simplified to 5.

### Serine hydrolase dendrogram

The serine hydrolase list was reported previously^67^. From this list, sequences were inputted into Geneious Prime (version 2020.1.2 Biomatters). Sequences were aligned using Clustal Omega, neighbor joining alignment. Dendrogram tree was generated using the Geneious Tree Builder; Genetic distance model Jukes-Cantor, Tree build method UPGMA, no outgroup, Bootstrap resampling, random seed 508,949, 300 interactions, support threshold of 1. CEST enzymes were colored red, PPA-UAR-1 was colored blue, and example proteases were colored green (Fig. S1).

## 2. Synthetic Procedures

**Synthesis of iglu#1 (15).** iglu#1 was synthesized as described previously^68^.

**Synthesis of angl#1 (17).** angl#1 was synthesized as described previously^69^.

Synthesis of 2-((*tert*-butoxycarbonyl)amino)benzoic acid (Boc-AA, SI-1).

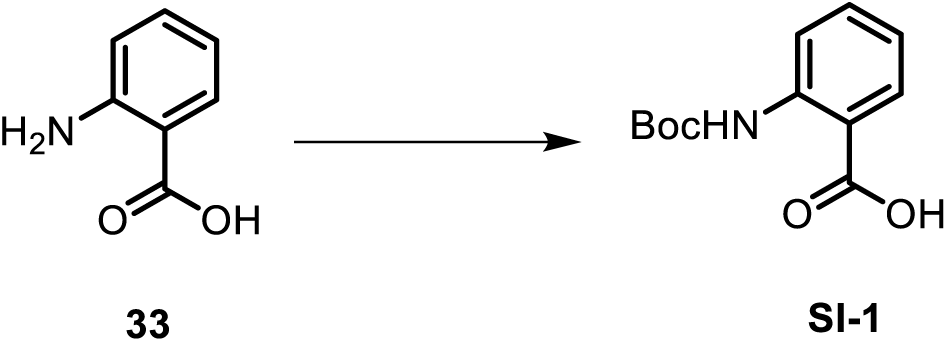

To a solution of anthranilic acid (**33**, 300 mg, 2.18 mmol) in 4 mL of THF and H_2_O (1:1), Boc-anhydride (520.8 mg, 2.39 mmol) was added, and 2 M NaOH was added to the mixture until pH 10 was reached. The reaction mixture was stirred at room temperature. After 23 hours, the solution was concentrated *in vacuo*, and 15% citric acid aqueous solution was added until pH 4 was reached. The white precipitate was filtered off and dried under vacuum to provide 2-((*tert*- butoxycarbonyl)amino)benzoic acid (**SI-1**, 496.7 mg, 96%) as a white solid. ^1^H NMR, 600 MHz, chloroform-d: δ (ppm) 10.06 (s, 1H), 8.47 (dd, J = 8.7, 0.9 Hz, 1H), 8.08 (dd, J = 7.9, 1.5 Hz, 1H), 7.57 (dt, J = 7.9, 1.5 Hz, 1H), 7.03 (dt, J = 7.2, 1.2 Hz, 1H), 1.55 (s, 9H).

### Synthesis of *N*-*β*-(6-(2ʹ-aminobenzoyl)-glucopyranosyl) indole (iglu#3, 34)

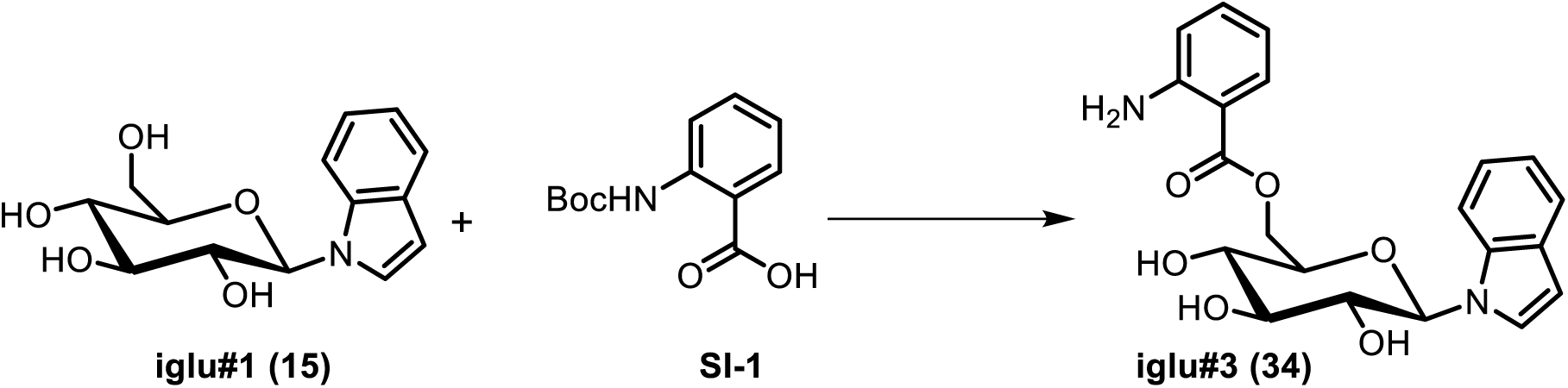

To a stirred solution of *N*-(*tert*-butoxycarbonyl)anthranilic acid^70^ (**SI-1**, 10 mg, 0.042 mmol) in dimethylformamide, 1-(3-dimethylaminopropyl)-3-ethylcarbodiimide hydrochloride (EDC·HCl, 20.1 mg, 0.105 mmol) was added. The mixture was stirred at room temperature for 5 min, and 4-dimethylaminopyridine (DMAP, 18.1 mg, 0.105 mmol) and *N*-*β*-glucopyranosyl indole (iglu#1, **15**, 9.8 mg, 0.0351 mmol) were added. The reaction mixture was stirred at room temperature. After 5 hours, the mixture was concentrated *in vacuo* to yield a viscous oil, which was dissolved in 1.4 mL of a 5:2 mixture of dichloromethane and methanol. Trifluoroacetic acid (TFA, 0.5 mL) was added slowly and the reaction mixture was stirred at room temperature. After 3 hours, the mixture was concentrated *in vacuo*. Preparative HPLC provided a pure sample of iglu#3 (**34**, 0.8 mg, 5.7%). See Table 5 for NMR spectroscopic data of iglu#3.

**Table 5.** NMR spectroscopic data for iglu#3 (34). ^1^H (600 MHz), HSQC, and HMBC NMR spectroscopic data were acquired in methanol-*d_4_*. Chemical shifts were referenced to δ(C**<u>H</u>**D_2_OD) = 3.31 ppm and δ(**^13^C**HD_2_OD) = 49.00 ppm.

| Position | $\delta^{13}\text{C}$ [ppm] | $\delta^1\text{H}$ [ppm] $J_{\text{HH}}$ [Hz] | HMBC |
| --- | --- | --- | --- |
| 1 | 86.9 | 5.51 ($J_{1,2} = 9.3$) | C-2, C-3, C-5, C-2', C-9' |
| 2 | 73.0 | 3.99 ($J_{2,3} = 9.0$) | C-1, C-3 |
| 3 | 78.7 | 3.65 ($J_{3,4} = 9.0$) | C-4 |
| 4 | 71.3 | 3.64 ($J_{4,5} = 9.1$) | C-3 |
| 5 | 77.5 | 3.91 ($J_{5,6a} = 5.5$) | C-4 |
| 6a | 64.1 | 4.43 ($J_{6a,6b} = 12.1$) | C-5, C-1'' |
| 6b | | 4.67 ($J_{5,6b} = 2.2$) | C-4, C-1'' |
| 2' | 126.3 | 7.37 ($J_{2',3'} = 3.3$) | C-1 (weak), C-3', C-4', C-8' (weak), C-9' |
| 3' | 102.9 | 6.48 |  |
| 4' | 130.4 |  |  |
| 5' | 121.4 | 7.52 ($J_{5',6'} = 8.0$) | C-3', C-7', C-9' |
| 6' | 120.8 | 7.03 ($J_{6',7'} = 7.4$,<br>$J_{3',6'} = 1.1$) | C-4', C-8' |
| 7' | 122.4 | 7.06 | C-5', C-9' |
| 8' | 111.5 | 7.53 | C-4', C-6' |
| 9' | 137.5 |  |  |
| 1'' | 168.6 |  |  |
| 2'' | 112.8 |  |  |
| 3'' | 132.1 | 7.90 ($J_{3'',4''} = 8.2$,<br>$J_{3'',5''} = 1.4$) | C-1'', C-5'', C-7'' |
| 4'' | 118.2 | 6.73 ($J_{4'',5''} = 7.6$) | C-2'', C-6'' |
| 5'' | 135.0 | 7.32 ($J_{5'',6''} = 7.8$) | C-3'', C-7'' |
| 6'' | 118.6 | 6.84 | C-2'', C-4'' |
| 7'' | 149.9 |  |  |

HRMS (ESI) *m/z*: [M - H]^-^ calcd for C_21_H_21_N_2_O_6_ 397.13938; found 397.14017.

### Synthesis of *N*-*β*-(6-nicotinoylglucopyranosyl) indole (iglu#5, SI-2)

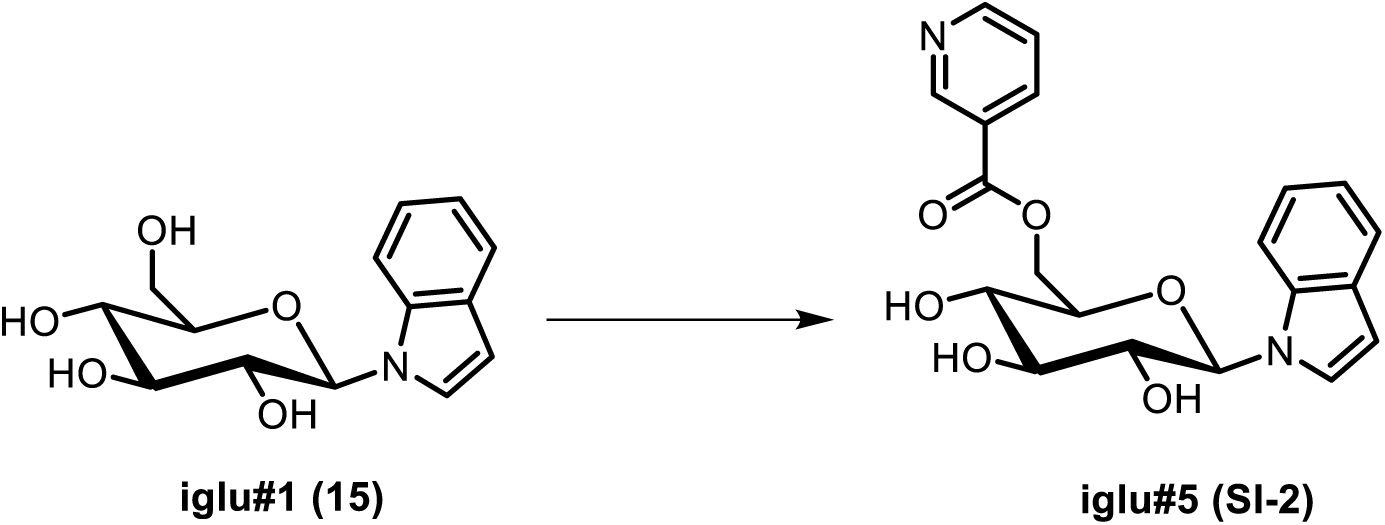

To a stirred solution of nicotinic acid (7.3 mg, 0.059 mmol) in a mixture of dimethylformamide and dichloromethane (1:1), EDC·HCl (28.4 mg, 0.148 mmol) was added. The mixture was stirred at room temperature for 30 min, before DMAP (18.1 mg, 0.148 mmol) and *N*-*β*- glucopyranosyl indole (iglu#1, **15**, 13.8 mg, 0.0494 mmol) were added. The reaction mixture was stirred at room temperature for 20 h, the mixture was concentrated *in vacuo*, and flash column chromatography on silica using a gradient of 0-25% methanol in dichloromethane afforded **iglu#5** (**SI-2**, 2.5 mg, 13.9%) as a colorless oil. See Table 6 for NMR spectroscopic data of iglu#5.

**Table 6.** NMR spectroscopic data for iglu#5 (SI-2). ^1^H (600 MHz), HSQC, and HMBC NMR spectroscopic data were acquired in methanol-*d_4_*. Chemical shifts were referenced to δ(C**<u>H</u>**D_2_OD) = 3.31 ppm and δ(**^13^C**HD_2_OD) = 49.00 ppm.

| Position | $\delta^{13}\text{C}$ ppm] | $\delta^1\text{H}$ [ppm] $J_{\text{HH}}$[Hz] | HMBC |
| --- | --- | --- | --- |
| 1 | 86.9 | 5.51 ($J_{1,2} = 9.2$) | C-2, C-3, C-5, C-2', C-9' |
| 2 | 73.0 | 4.00 ($J_{2,3} = 9.0$) | C-1, C-3 |
| 3 | 78.7 | 3.65 ($J_{3,4} = 9.0$) | C-4 |
| 4 | 71.4 | 3.63 ($J_{4,5} = 8.9$) | C-3 |
| 5 | 77.4 | 3.95 ($J_{5,6a} = 5.8$) | C-4 |
| 6a | 65.3 | 4.51 ($J_{6a,6b} = 12.1$) | C-4, C-5, C-1'' |
| 6b | | 4.75 ($J_{5,6b} = 2.3$) | C-4, C-5, C-1'' |
| 2' | 126.4 | 7.37 ($J_{2',3'} = 3.5$) | C-3', C-4', C-9' |
| 3' | 103.1 | 6.47 | C-2', C-4', C-9' |
| 4' | 130.5 |  |  |
| 5' | 121.4 | 7.51 ($J_{5',6'} = 7.9$) | C-4', C-6', C-9' |
| 6' | 120.8 | 7.01 ($J_{6',7'} = 7.5$,<br>$J_{3',6'} = 1.2$) | C-4', C-8' |
| 7' | 122.5 | 7.05 | C-4', C-5', C-8', C-9' |
| 8' | 111.4 | 7.49 | C-4', C-6' |
| 9' | 137.6 |  |  |
| 1'' | 165.8 |  |  |
| 2'' | 127.7 |  |  |
| 3'' | 150.8 | 9.12 ($J_{3'',6''} = 0.5$,<br>$J_{3'',7''} = 2.0$) | C-2'', C-5'', C-7'' |
| 5'' | 153.7 | 8.74 ($J_{5'',6''} = 4.9$,<br>$J_{5'',7''} = 1.7$) | C-3'', C-6'', C-7'' |
| 6'' | 125.1 | 7.54 ($J_{6'',7''} = 8.0$) | C-2'', C-5'' |
| 7'' | 138.9 | 8.37 | C-1'', C-2'', C-5'' |

HRMS (ESI) *m/z*: [M + H]^+^ calcd for C_20_H_21_N_2_O_6_ 385.13941; found 385.14038.

### Synthesis of *N*-*β*-(6-(2ʹ-methylbut-2ʹ*E*-enoyl)-glucopyranosyl) indole (iglu#7, SI-3)

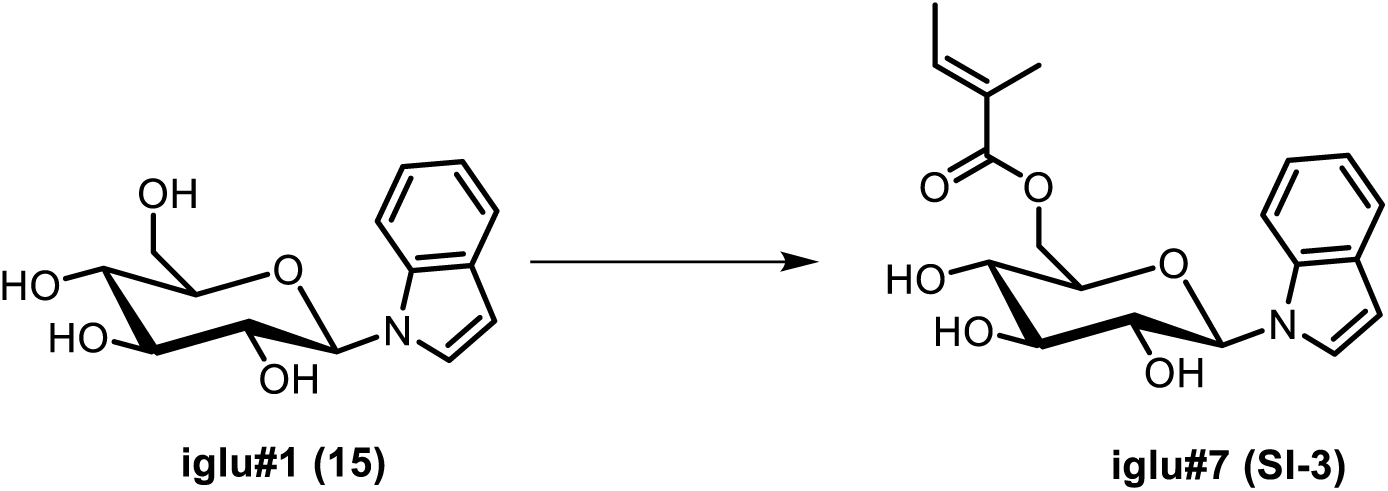

To a stirred solution of tiglic acid (5.0 mg, 0.050 mmol) in a 1:1 mixture of dimethylformamide and dichloromethane, EDC·HCl (23.9 mg, 0.125 mmol) was added. The mixture was stirred at room temperature for 30 min, and DMAP (15.2 mg, 0.125 mmol) and *N*-*β*-glucopyranosyl indole (iglu#1, **15**, 11.6 mg, 0.0416 mmol) were added. The reaction mixture was stirred at room temperature for 22 hours and then concentrated *in vacuo*. Flash column chromatography on silica using a gradient of 0-30% methanol in dichloromethane afforded **iglu#7** (**SI-3**, 2.5 mg, 11.3%) as a colorless oil. See Table 7 for NMR spectroscopic data of iglu#7.

**Table 7.** NMR spectroscopic data for iglu#7 (SI-3). ^1^H (600 MHz), HSQC, and HMBC NMR spectroscopic data were acquired in methanol-*d_4_*. Chemical shifts were referenced to δ(C**<u>H</u>**D_2_OD) = 3.31 ppm and δ(**^13^C**HD_2_OD) = 49.00 ppm.

| Position | $\delta^{13}\text{C}$ ppm] | $\delta^1\text{H}$ [ppm] $J_{\text{HH}}[\text{Hz}]$ | HMBC |
| --- | --- | --- | --- |
| 1 | 86.9 | 5.46 ($J_{1,2} = 9.1$) | C-2, C-3, C-5, C-2', C-9' |
| 2 | 73.2 | 3.96 ($J_{2,3} = 9.0$) | C-1, C-3 |
| 3 | 78.9 | 3.61 ($J_{3,4} = 9.0$) | C-2, C-4 |
| 4 | 71.4 | 3.55 ($J_{4,5} = 9.6$) | C-3, C-5, C-6 |
| 5 | 77.6 | 3.81 ($J_{5,6a} = 5.6$) | C-1 (weak), C-3, C-4 |
| 6a | 64.5 | 4.27 ($J_{6a,6b} = 11.9$) | C-4, C-5, C-1'' |
| 6b | | 4.49 ($J_{5,6b} = 2.2$) | C-4, C-5, C-1'' |
| 2' | 126.6 | 7.35 ($J_{2',3'} = 3.5$) | C-1 (weak), C-3', C-4', C-5' (weak), C-8' (weak), C-9' |
| 3' | 103.2 | 6.48 |  |
| 4' | 130.6 |  |  |
| 5' | 121.6 | 7.53 ($J_{5',6'} = 7.9$) | C-3', C-7', C-9' |
| 6' | 120.9 | 7.05 ($J_{6',7'} = 7.5$,<br>$J_{3',6'} = 1.1$) | C-4', C-8', C-9' (weak) |
| 7' | 122.5 | 7.11 | C-5', C-8' (weak), C-9' |
| 8' | 111.7 | 7.50 | C-4', C-6' |
| 9' | 137.6 |  |  |
| 1'' | 169.2 |  |  |
| 2'' | 129.3 |  |  |
| 3'' | 138.9 | 6.87 ($J_{3'',4''} = 6.8$) | C-1'', C-4'', C-5'' |
| 4'' | 14.2 | 1.79 | C-2'', C-3'' |
| 5'' | 11.9 | 1.81 | C-1'', C-2'', C-3'' |

HRMS (ESI) *m/z*: [M + H]^+^ calcd for C_19_H_24_NO_6_ 362.15981; found 362.16025.

### Synthesis of *N*-*β*-(6-(pyrrole-2ʹ-carbonyl)-glucopyranosyl) indole (iglu#9, SI-4)

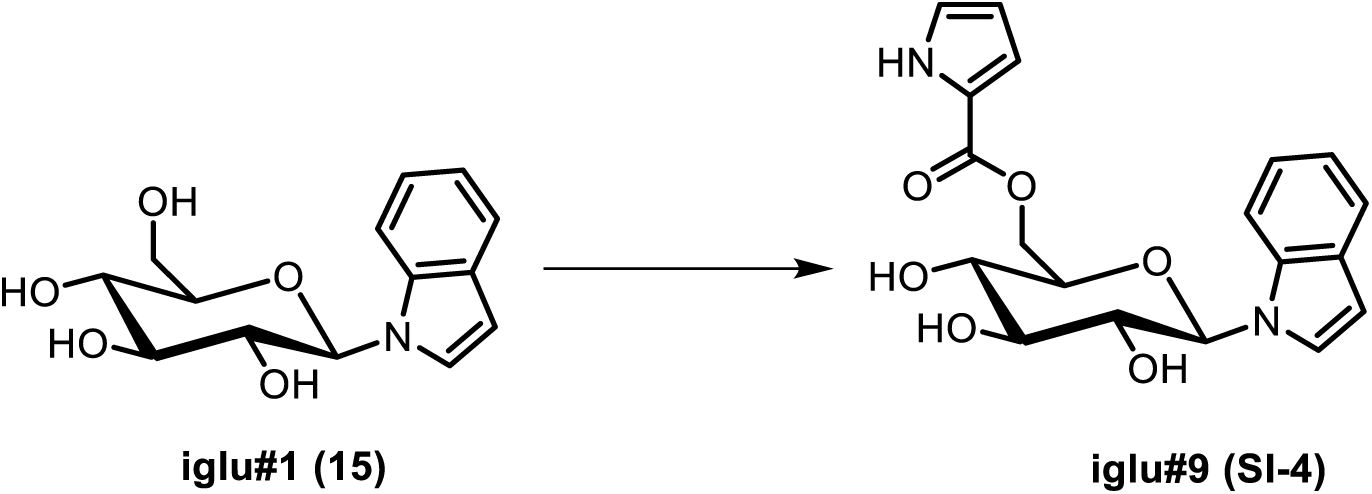

To a suspension of pyrrole-2-carboxylic acid (6.0 mg, 0.054 mmol) in dichloromethane, oxalyl chloride (14 µL, 0.163 mmol) was added slowly, followed by dimethylformamide (1 µL, 0.0129 mmol). The mixture was stirred at room temperature for 18 hours and then concentrated to dryness *in vacuo*. The residue was re-dissolved in dimethylformamide (2 mL) containing *N*-*β*- glucopyranosyl indole (iglu#1, **15**, 10.8 mg, 0.0387 mmol). Triethylamine (45 µL, 0.324 mmol) was added, and the reaction was stirred at 35 °C for 7 days. Subsequently the mixture was concentrated *in vacuo*, and flash column chromatography on silica using a gradient of 0-30% methanol in dimethylformamide afforded **iglu#9** (**SI-4**, 1.5 mg, 10.4%) as a colorless oil. See Table 8 for NMR spectroscopic data of iglu#9.

**Table 8.** NMR spectroscopic data for iglu#9 (SI-4). ^1^H (600 MHz), HSQC, and HMBC NMR spectroscopic data were acquired in methanol-*d_4_*. Chemical shifts were referenced to δ(C**<u>H</u>**D_2_OD) = 3.31 ppm and δ(**^13^C**HD_2_OD) = 49.00 ppm.

| Position | $\delta^{13}\text{C}$ [ppm] | $\delta^1\text{H}$ [ppm] $J_{\text{HH}}$ [Hz] | HMBC |
| --- | --- | --- | --- |
| 1 | 86.9 | 5.47 ($J_{1,2} = 9.1$) | C-2, C-3, C-5, C-2', C-9' |
| 2 | 73.2 | 3.96 ($J_{2,3} = 9.0$) | C-1, C-3 |
| 3 | 78.7 | 3.62 ($J_{3,4} = 9.8$) | C-4 |
| 4 | 71.3 | 3.61 ($J_{4,5} = 9.7$) | C-3 |
| 5 | 77.9 | 3.86 ($J_{5,6a} = 5.7$) | |
| 6a | 63.9 | 4.38 ($J_{6a,6b} = 11.9$) | C-5, C-1'' |
| 6b | | 4.68 ($J_{5,6b} = 2.1$) | C-4, C-1'' |
| 2' | 126.6 | 7.36 ($J_{2',3'} = 3.4$) | C-3', C-4', C-9' |
| 3' | 103.1 | 6.47 | C-2', C-4', C-9' |
| 4' | 130.6 |  |  |
| 5' | 121.4 | 7.52 ($J_{5',6'} = 7.8$) | C-7', C-9' |
| 6' | 120.8 | 7.02 ($J_{6',7'} = 7.3$,<br>$J_{3',6'} = 1.2$) | C-4', C-8' |
| 7' | 122.4 | 7.05 | C-5', C-9' |
| 8' | 111.6 | 7.50 | C-4', C-6' |
| 9' | 137.4 |  |  |
| 1'' | 162.4 |  |  |
| 2'' | 123.0 |  |  |
| 4'' | 124.7 | 6.96 ($J_{4'',5''} = 2.5$,<br>$J_{4'',6''} = 1.4$) | C-2'', C-5'', C-6'' |
| 5'' | 110.6 | 6.20 ($J_{5'',6''} = 3.8$) | C-2''(weak), C-4''(weak) |
| 6'' | 116.8 | 6.90 | C-2'', C-4'', C-5'' |

**Table 9.**
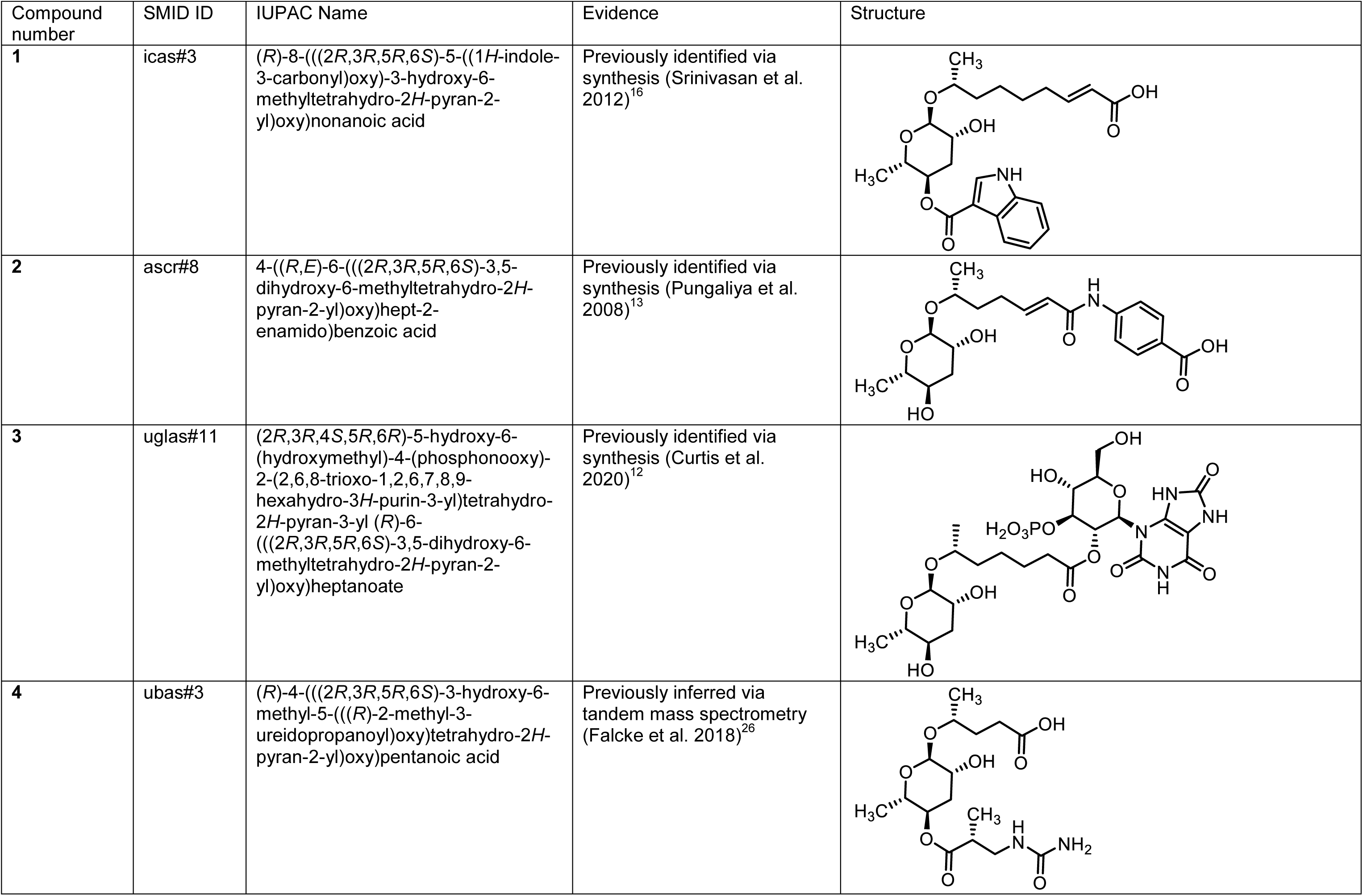
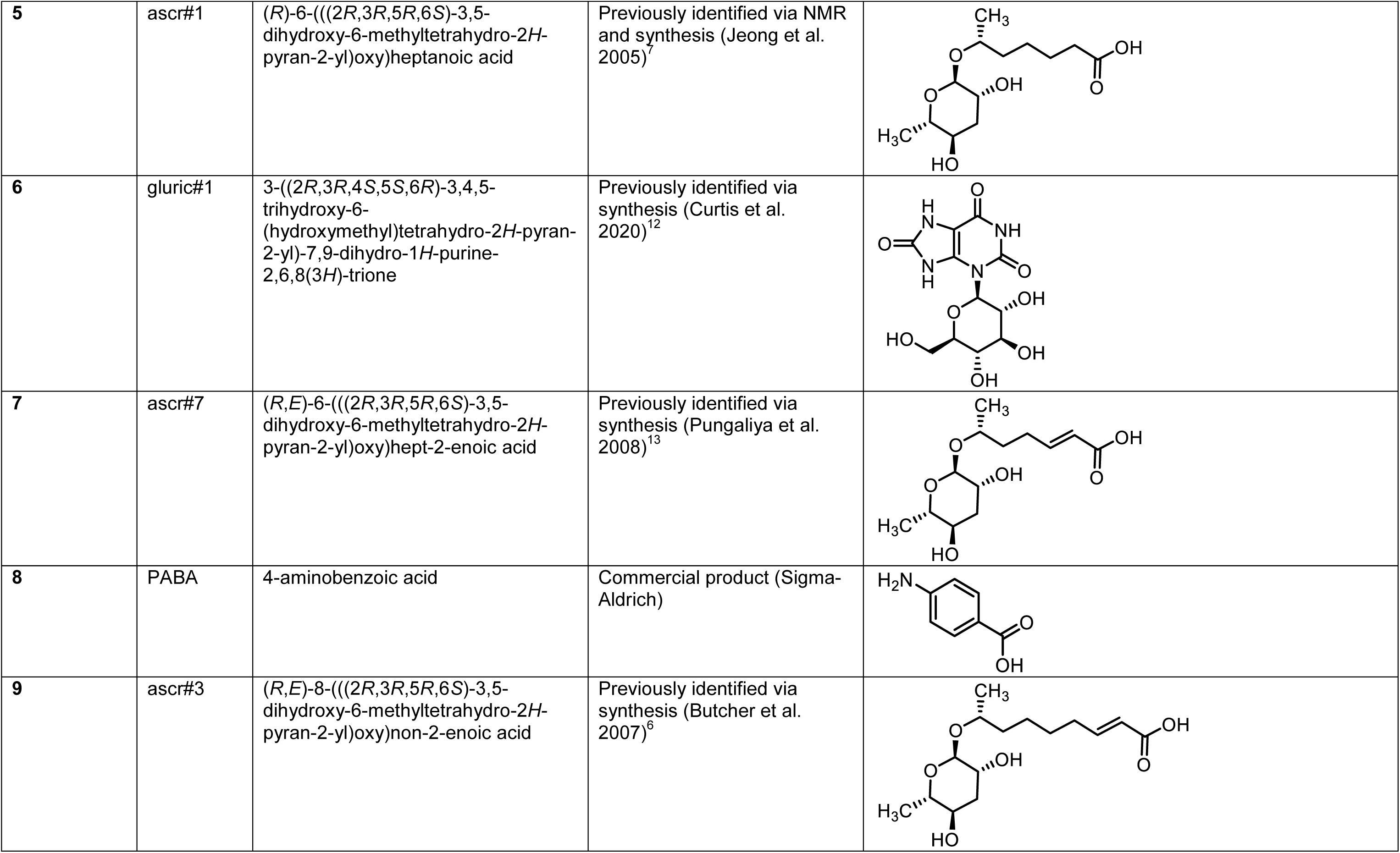
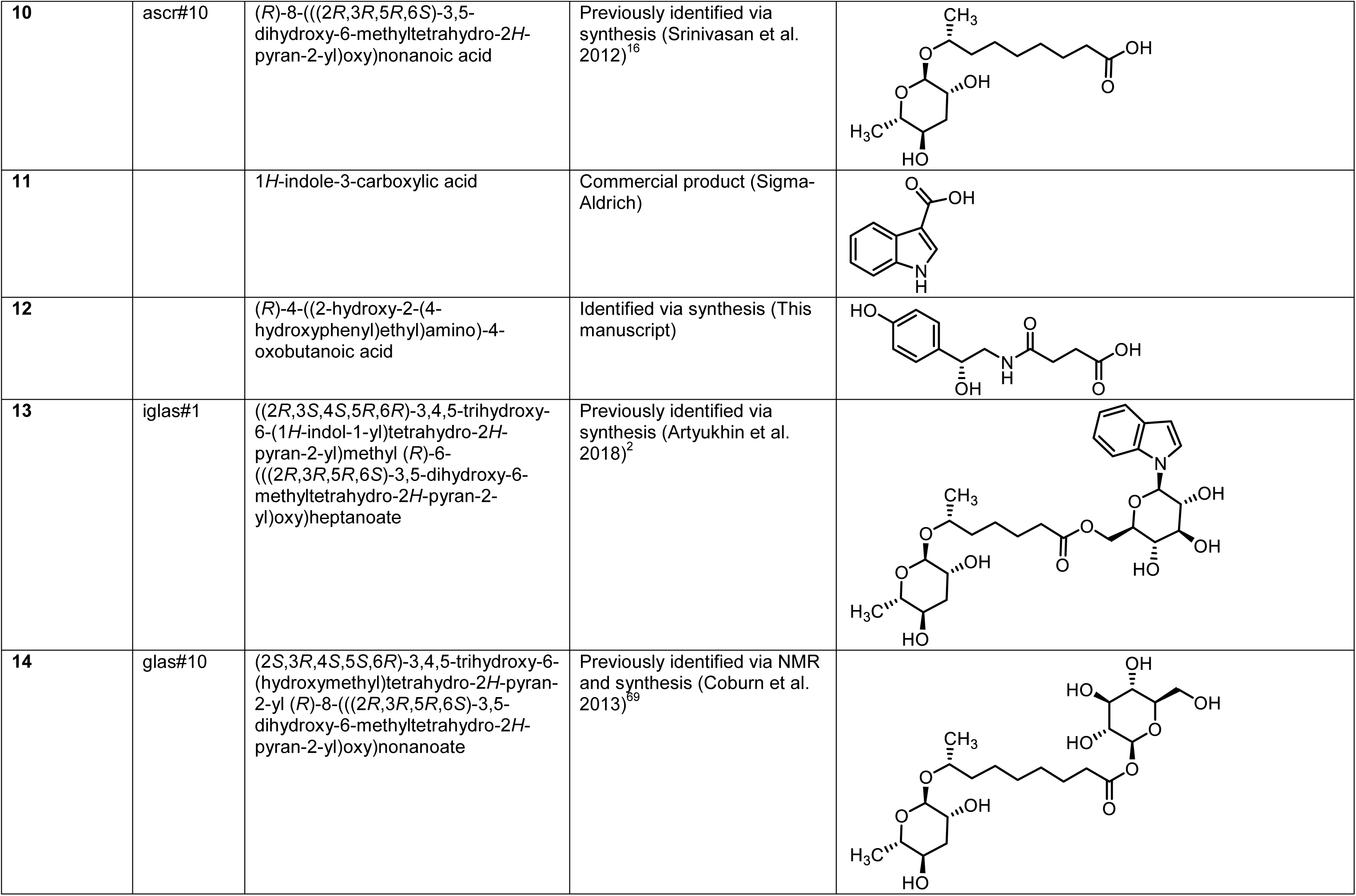
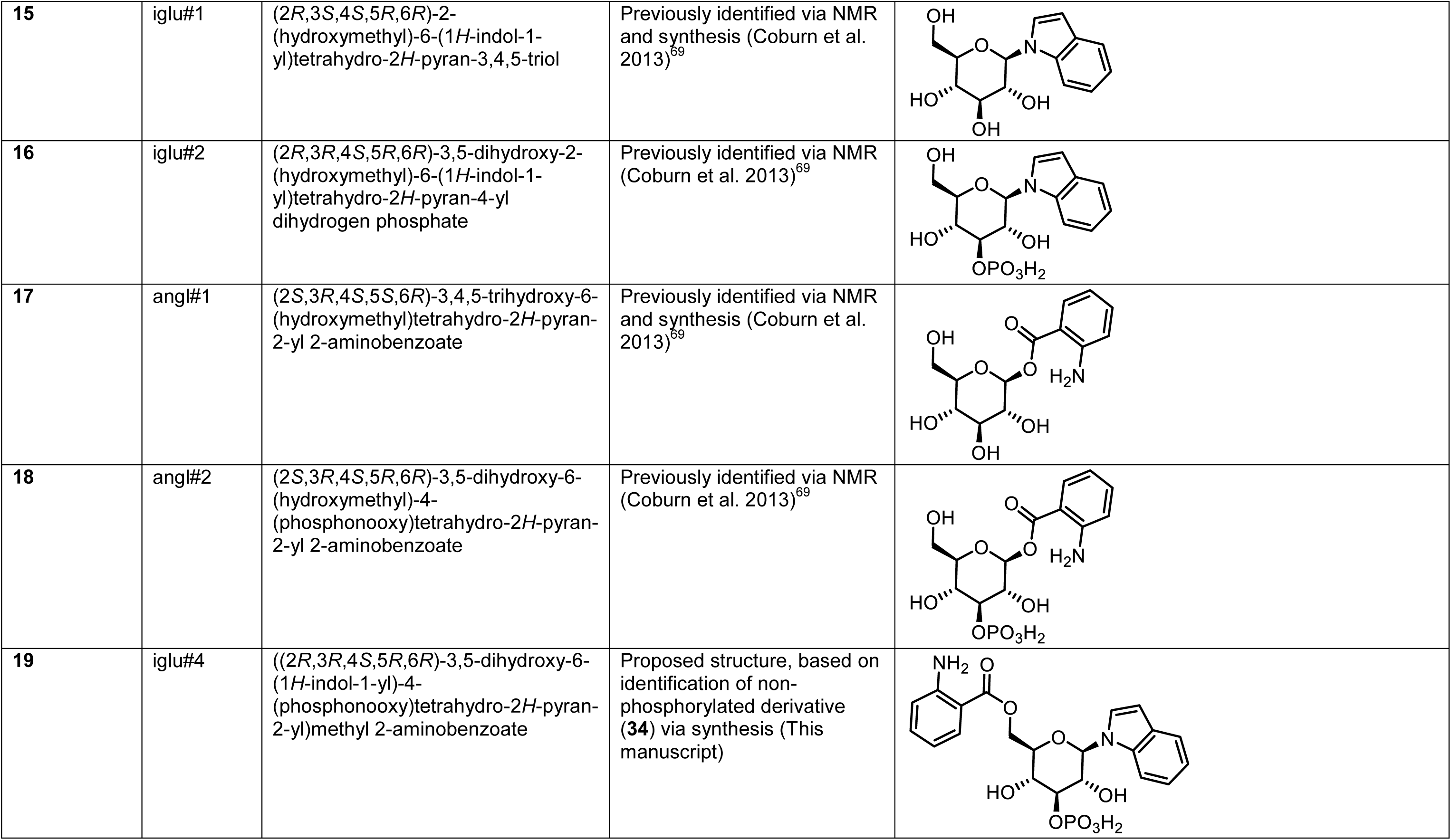
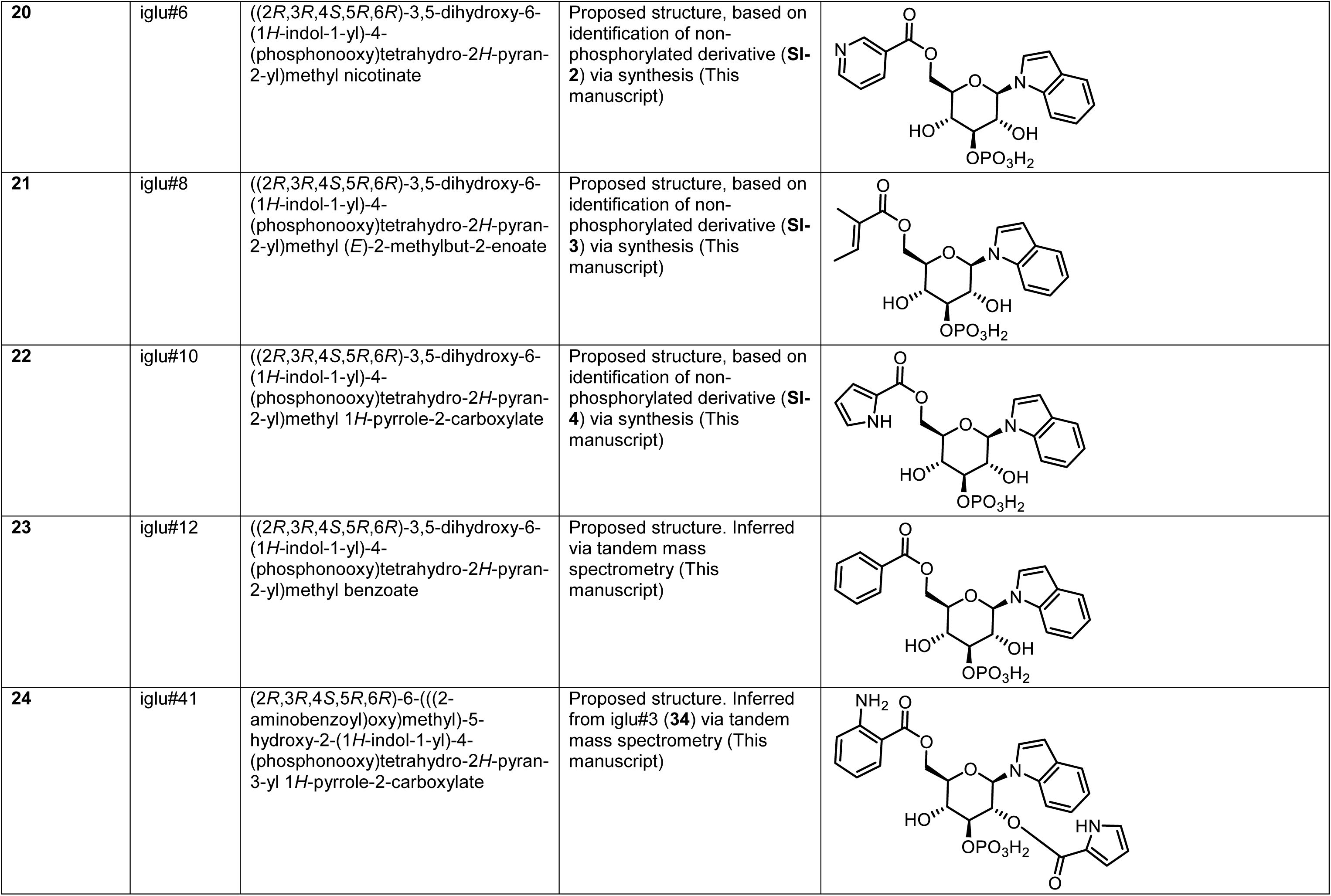
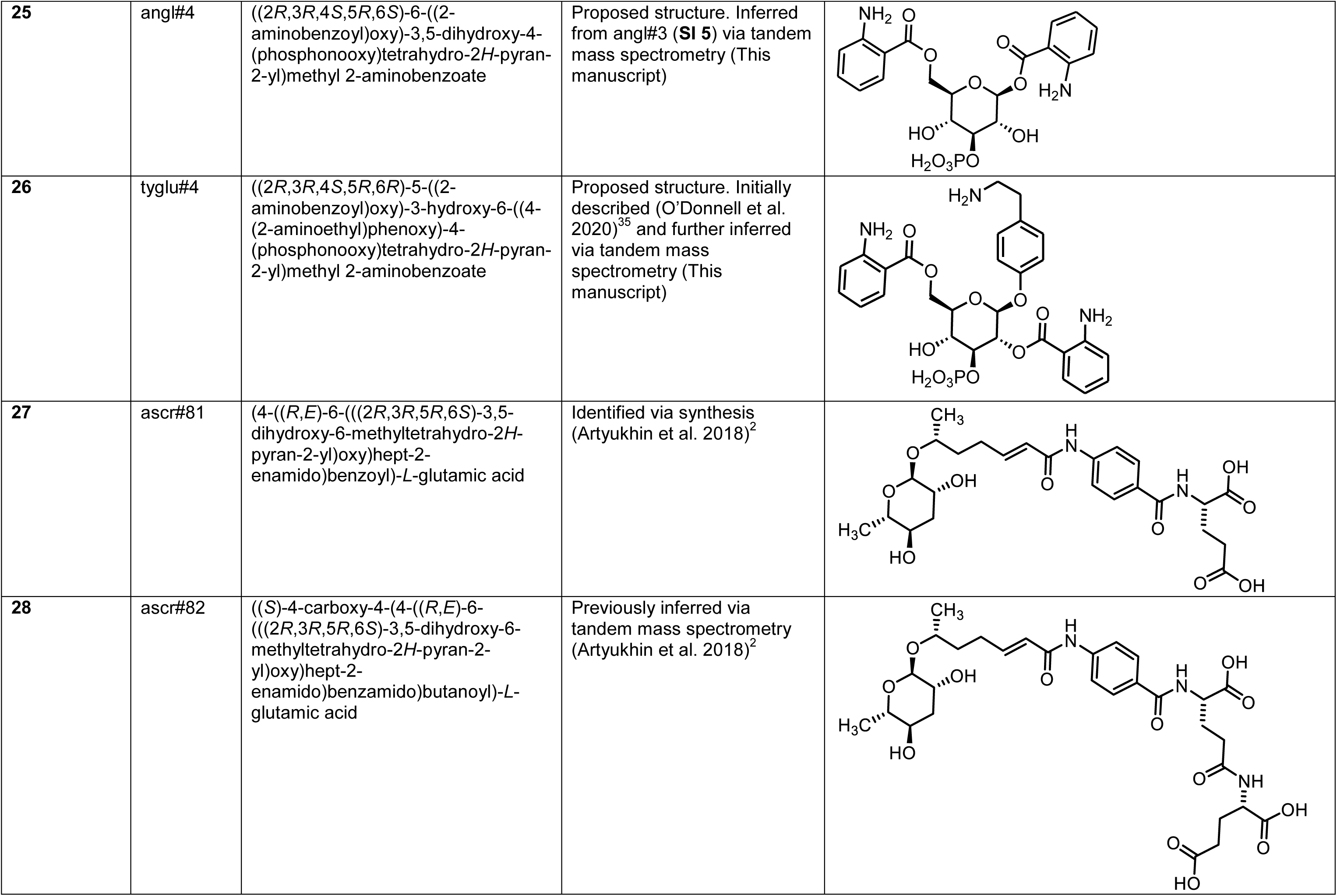
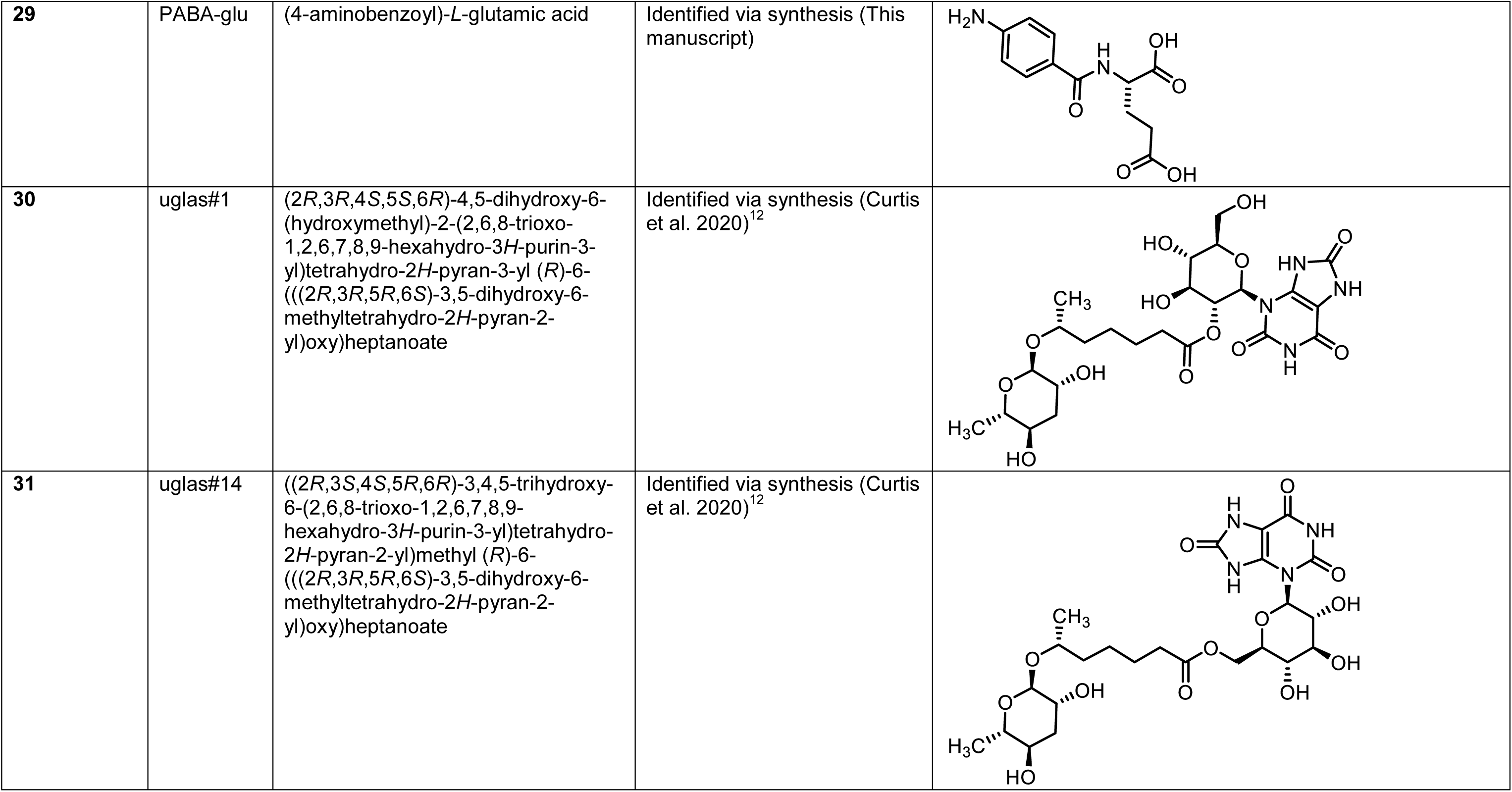
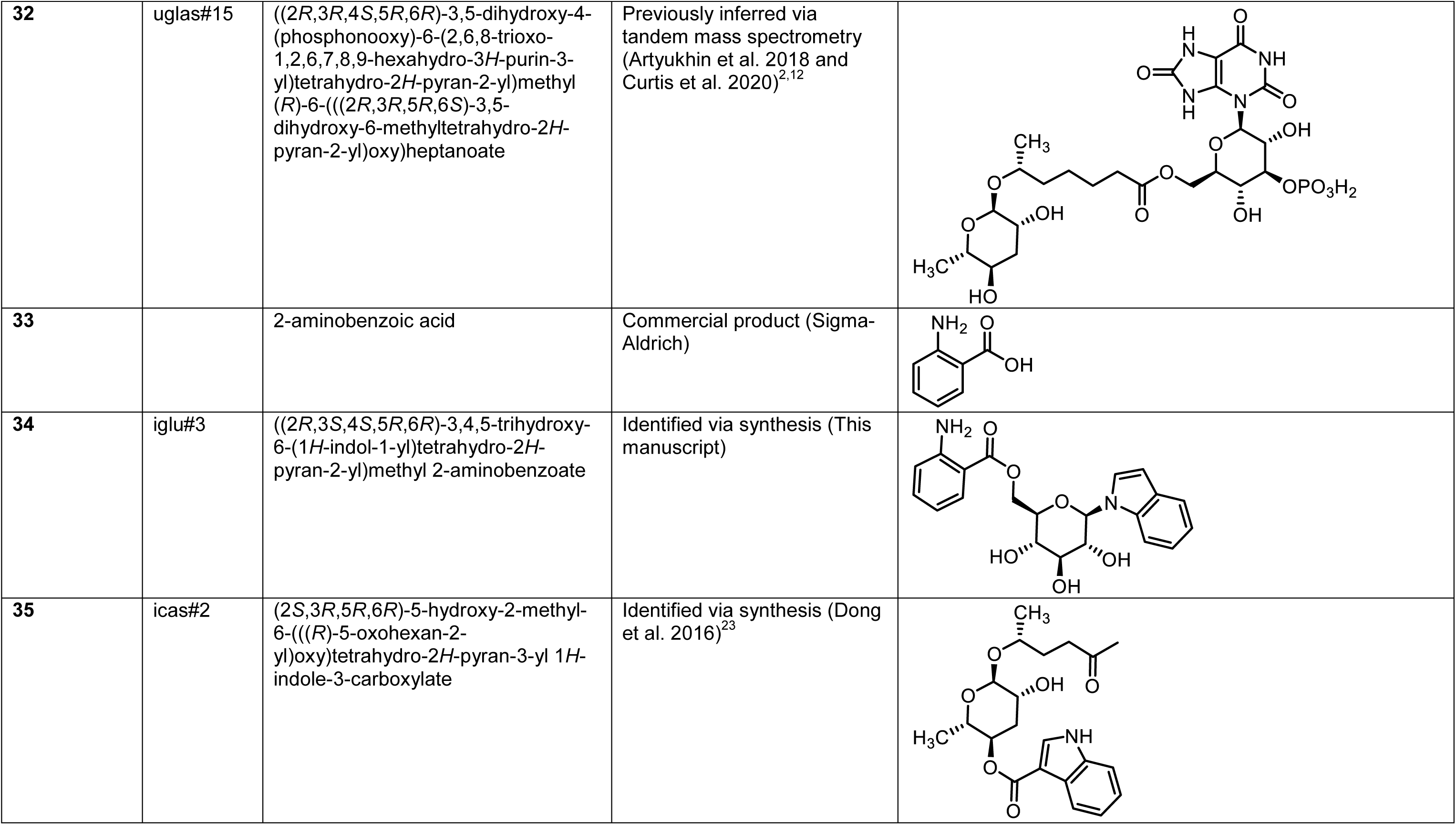
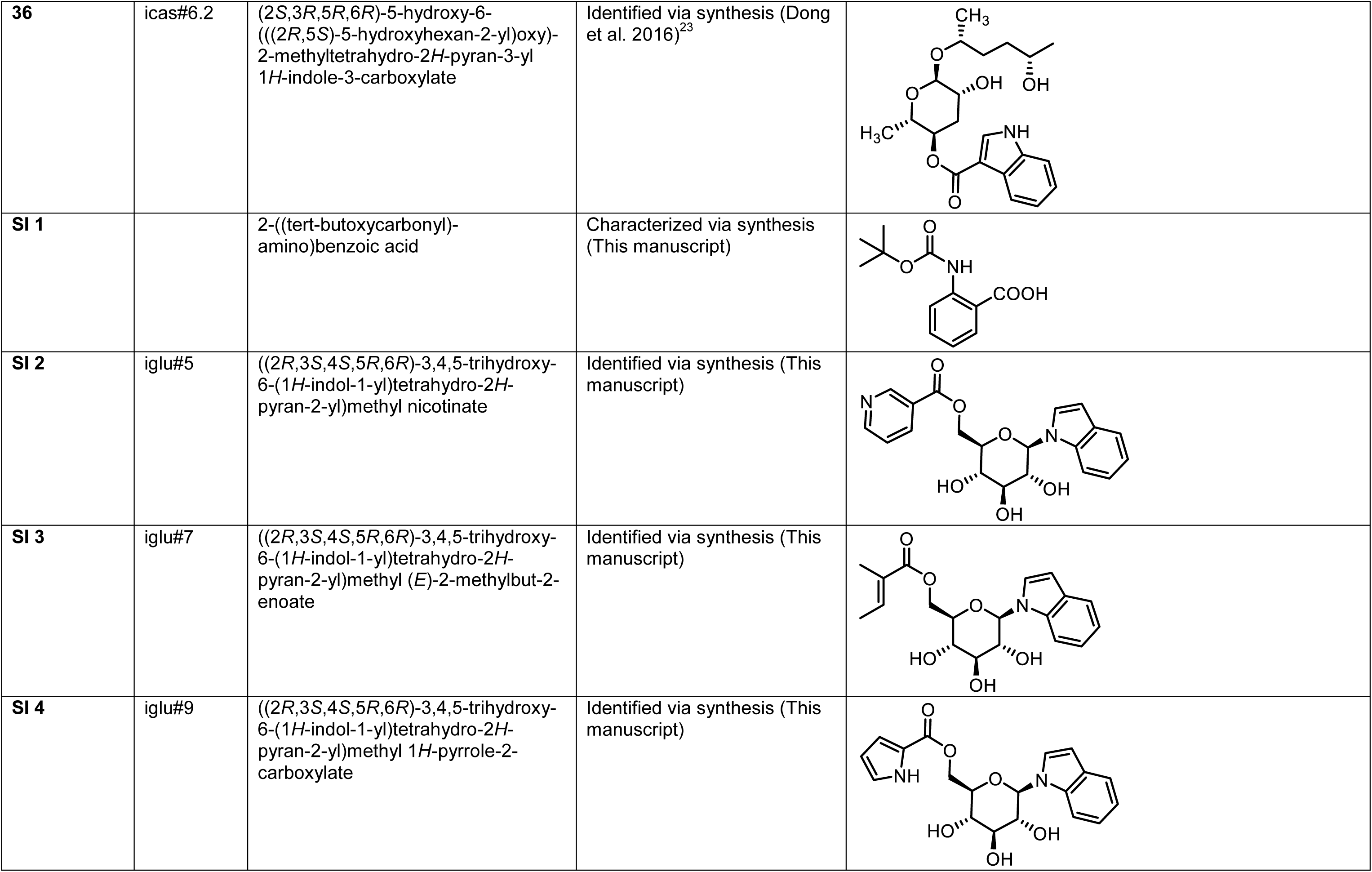
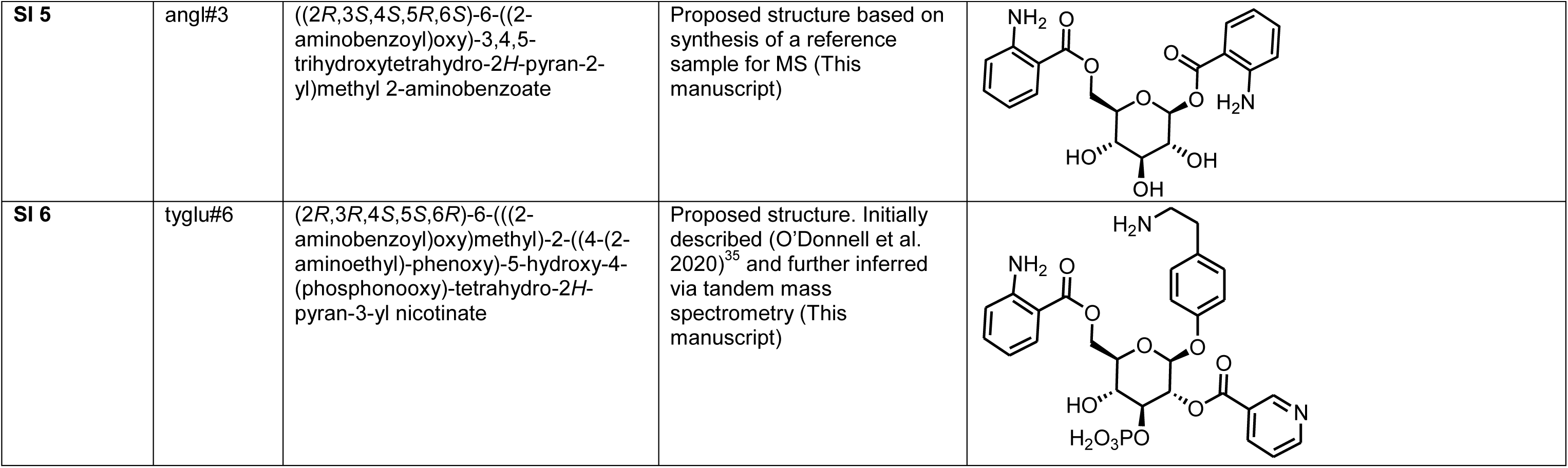
List of all modular metabolites referred to in the text and Figures.

**Table 10.** Compiled data depicted in Figures 1, 3, 4, S6, S7, S9, S10, S11, S13, and S16. Attached as a separate file.

HRMS (ESI) *m/z*: [M + H]^+^ calcd for C_19_H_21_N_2_O_6_ 373.13941; found 373.14026.

### Synthesis of an HPLC standard of ((2*R*,3*S*,4*S*,5*R*,6*S*)-6-((2-aminobenzoyl)oxy)-3,4,5- trihydroxytetrahydro-2*H*-pyran-2-yl)methyl 2-aminobenzoate (angl#3, SI-5)

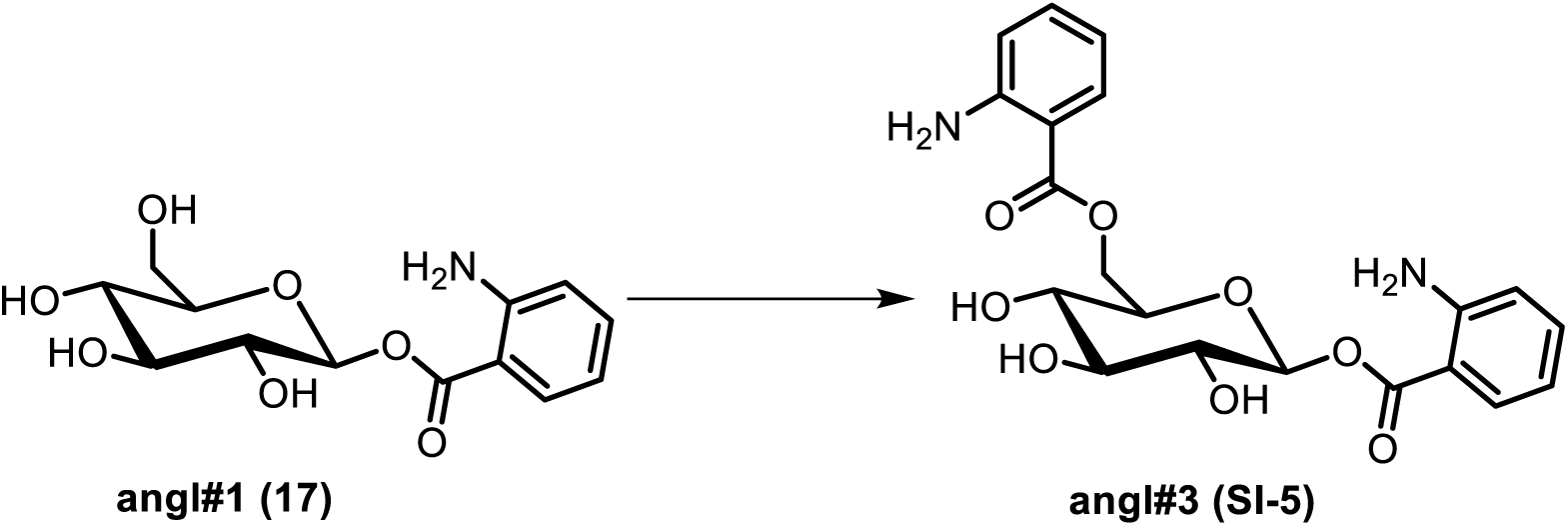

To a stirred solution of Boc-AA (2 mg, 0.00 84mmol) in dimethylformamide, 1-(3- dimethylaminopropyl)-3-ethylcarbodiimide hydrochloride (EDC·HCl) (3.9 mg, 0.0203 mmol) was added. The mixture was stirred at room temperature for 5 min, and 4-dimethylaminopyridine (DMAP) (2.5 mg, 0.0203 mmol) and angl#1 (**17**, 2 mg, 0.0068 mmol) were added. The reaction mixture was stirred at room temperature. After 5 hours, the mixture was concentrated *in vacuo*. The crude product was dissolved in 0.55 mL dichloromethane and methanol (10:1) and trifluoroacetic acid (TFA, 500 µL) was added slowly. The reaction mixture was stirred at room temperature for 3 hours then was concentrated *in vacuo*, affording **angl#3** (**SI-5**).

HRMS (ESI) *m/z*: [M + H]^+^ calcd for C_20_H_23_N_2_O_7_ 403.14998; found 403.15100.

### Synthesis of an HPLC standard of *N*-(*p*-aminobenzoyl)glutamate (PABA-glutamate) (29)

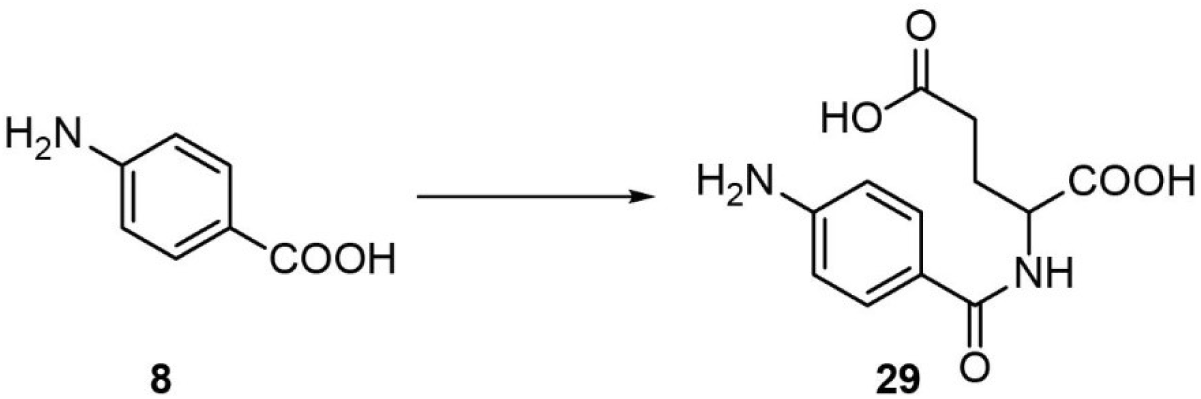

*p*-Aminobenzoic acid (Chem-Impex) (**8**) was dissolved in warm dichloromethane (DCM) containing triethylamine (0.1 eq). EDC·HCl (Amresco Biochemicals) (1 eq) and di-*tert*-butyl glutamate (1 eq) was added to the reaction mixture. *N*,*N*-Dimethylaminopyridine (1.1 eq) was then added to the mixture and the resulting mixture was stirred at room temperature for 24 hr and then extracted with ethyl acetate. The organic layer was dried with sodium sulfate and evaporated to dryness *in vacuo*. The crude product was dissolved in DCM, and trifluoroacetic acid (TFA) was added (100 eq). The reaction was then stirred for 6 hr at room temperature. TFA and DCM were evaporated off to yield crude PABA-glutamate (**29**). ^1^H NMR, 600 MHz, methanol: δ (ppm) 7.93 (d, 8.6 Hz, 2H), 7.37 (d, 8.5 Hz, 2H), 4.61 (dd, 5.0, 9.3 Hz, 1H), 2.09-2.28 (m, 4H).

## Author Contributions

The manuscript was written through contributions of all authors and all authors have given approval to the final version of the manuscript.

^‡^These authors contributed equally.

## Acknowledgements

This research was funded by an NIH Chemical Biology Interface (CBI) Training Grant 5T32GM008500 (to B.C.), National Institutes of Health grants R35 GM131877 (to F.C.S.), and R24OD023041 (to P.W.S.). F.C.S. is a Faculty Scholar of the Howard Hughes Medical Institute. We thank WormBase for sequences, Tsui-Fen Chou for Cas9 protein, Ying (Kitty) Zhang for assistance with NMR spectroscopy, and Navid Movahed for assistance with mass spectrometry.

